# Aggressive Cholesterol Lowering Normalizes Atherosclerosis Regression in *Jak2^V617F^* Mice

**DOI:** 10.1101/2025.07.23.666334

**Authors:** Brian D. Hardaway, Trevor P. Fidler, Mojdeh Tavallaie, Kleopatra Avrampou, Cheng-Chieh Hsu, Sandra Schiavone, Tong Xiao, Nan Wang, Alan R. Tall

## Abstract

**Background:** The *Jak2^V617F^* (*Jak2^VF^*) mutation is an important cause of both clonal hematopoiesis of indeterminate potential (CHIP) and myeloproliferative neoplasms (MPN). Mouse models of *Jak2^VF^* CHIP and MPN show accelerated atherosclerosis progression, driven by macrophage inflammasome activation. We undertook the present study to assess the hypothesis that ongoing inflammation would impede atherosclerosis resolution in *Jak2^VF^* mice.

**Methods and Results:** Chimeric *Jak2^VF/WT^* or control *WT/WT* bone marrow was transplanted into *Ldlr^−/-^*mice and, following 13-16 weeks of Western diet-induced atherosclerosis progression, cholesterol was lowered either moderately (to 200-300 mg/dl) or markedly (to 100 mg/dl). With moderate cholesterol lowering, there was impaired resolution of lesions in *Jak2^VF^*MPN mice compared to controls. However, with marked cholesterol lowering, progression of lesions was halted in both *Jak2^VF^* MPN and control mice while macrophage burden was decreased and lesional collagen was increased similarly in *Jak2^VF^*MPN and control mice.

Two mechanisms of low-density lipoprotein (LDL) lowering-induced suppression of inflammation in plaques were implicated: 1) reversal of increased proliferation, DNA damage and Absent in Melanoma 2 (AIM2) inflammasome activation specifically in *Jak2^VF^* macrophages and 2) markedly increased macrophage triggering receptor expressed on myeloid cells 2 (TREM2), c-myc expressing macrophages in both *Jak2^VF^* and control mice.

**Conclusions:** Aggressive LDL lowering reverses inflammasome activation and induces pro-resolving changes in macrophages in *Jak2^VF^* MPN, halting atherosclerosis progression and promoting features of plaque stabilization. These findings suggest that aggressive LDL cholesterol lowering could reverse atherosclerotic cardiovascular disease (ACVD) risk in individuals with *JAK2^VF^*CHIP or MPN.

## 1. Introduction

Despite therapeutic advances, atherosclerotic cardiovascular disease (ACVD) remains the leading cause of death in the developed world (1). Although inflammation plays a key role in atherosclerosis progression, cholesterol accumulation remains a prerequisite for lesion development, and low-density lipoprotein (LDL) lowering has consistently shown a benefit in clinical trials (2,3,4).

The role of inflammation in increasing ACVD risk has also been well established. The LoDoCo 2 trial found that the risk of cardiovascular events was reduced in patients with chronic coronary disease treated with colchicine (5). Based on these and other results, the U.S. Food and Drug Administration approved the use of low dose colchicine as the first anti-inflammatory drug for heart disease (6). The CANTOS trial involving the use of interleukin-1β (IL-1β) antibodies showed reduced cardiovascular events in high-risk patients with mean LDL cholesterol levels of 81 mg/dl - 84 mg/dl and a high-sensitivity C-reactive protein (hs-CRP) level of at least 2 mg/L. However, there was a small increase in fatal infections, and the FDA did not approve this treatment for marketing (7). Many observations suggest that hypercholesterolemia is linked to inflammatory risk. For example, statins lower hs-CRP levels (8) and preclinical studies have shown that cholesterol accumulation in plaque macrophages promotes inflammasome activation (9,10,11,12). However, the mechanisms linking hypercholesterolemia, macrophage cholesterol accumulation and inflammation remain incompletely understood, especially in the clinically important context of atherosclerosis regression induced by LDL lowering (13,14,15,16,17,18,19).

Clonal hematopoiesis (CH) arises from leukemogenic mutations in hematopoietic stem cells (HSC) that confer a selective advantage and lead to clonal expansion of blood cells. CH commonly arises from mutations in genes that mediate epigenetic modifications or cytokine signaling (20,21). Clonal hematopoiesis of indeterminate potential (CHIP) is defined as having a mutant allele burden >2% in blood cells in the absence of changes associated with hematopoietic malignancy (20). Amongst CHIP mutations, *JAK2^VF^*, which increases signaling by hematopoietic cytokines, confers the greatest risk of ACVD (22,23,24). Moreover, *JAK2^VF^* has been detected in 3-4% of a general European population (25). *JAK2^VF^* CHIP and myeloproliferative neoplasms (MPN) appear to form a disease spectrum, with both increasing atherosclerotic and thrombotic risk (26,27). Similarly, both *Jak2^VF^* MPN and low allele burden *Jak2^VF^*CHIP in mice that lack changes in blood cell counts or splenomegaly increase thrombosis and atherosclerosis progression involving leukocyte mediated inflammatory mechanisms (28,29). Additionally, we showed in a hypercholesterolemic mouse model that *Jak2^VF^* CHIP or MPN exacerbates atherosclerosis via inflammasome activation (30) and genetic studies suggest a similar role in humans (31). Based on these findings, we hypothesized that LDL lowering-induced changes in plaques such as halted progression and beneficial remodeling would be impaired in *Jak2^VF^* mice due to persistence of the underlying mechanisms promoting inflammasome activation. Contrary to our hypothesis, we found that cholesterol lowering reversed inflammasome associated changes in *Jak2^VF^* macrophages and that marked cholesterol lowering induced beneficial lesional changes similarly in *Jak2^VF^* MPN mice and controls. These studies suggest the importance of rigorous control of LDL cholesterol levels in individuals with *JAK2^VF^*CHIP or MPN.

## 2. Methods

Several experimental procedures were performed as previously described (16,30).

### 2.1 Anesthesia and Euthanasia

Mice were anesthetized using 5% isoflurane via inhalation at a flow rate of approximately 3L/min oxygen using a calibrated vaporizer. Anesthetic depth was assessed using respiratory rate monitoring and the toe-pinch reflex. Isoflurane was administered acutely during procedures requiring anesthesia, including blood collection and euthanasia. Blood collection was performed via retro-orbital sinus for plasma, complete blood counts, and flow cytometric analysis at the indicated timepoints. For euthanasia, mice were first rendered deeply unconscious with isoflurane, after which cervical dislocation was performed as a secondary method to ensure death, in accordance with institutional IACUC approval, NIH guidelines, and the AVMA Guidelines for the Euthanasia of Animals (2020 Edition).

### 2.2 Mice

All mice used for these studies were male and on a C57BL/6J background. Mice were housed in a pathogen-free facility under standard conditions of temperature (23^°^C) with a 12-hour light-dark with ad libitum food and water access. Cages and water were changed every 7-14 days. All mouse experiments were approved by the Institutional Animal Care and Use Committee of Columbia University under protocol number AABG0561 and were conducted in accordance with NIH guidelines and the Guide for the Care and Use of Laboratory Animals. Mice heterozygous for the *Jak2^VF^*conditional knock-in allele were generated as previously described (32). *Jak2^VF^* mice were crossed to *Mx1-Cre* (The Jackson Laboratory, B6.Cg-Tg(*Mx1-Cre*)1*Cgn*/J (003556) or *Scl-Cre* (The Jackson Laboratory, C57BL/6-Tg(Tal1-cre/ERT)42-056Jrg/J (037466) to generate *Mx1-CreJak2^VF^* and *Scl-CreJak2^VF^* mice respectively. CD45.1*Mx1-CreJak2^VF^* mice were generated by crossing *Mx1-CreJak2^VF^* mice to CD45.1 (The Jackson Laboratory B6.SJL-Ptprca Pepcb/BoyJ (002014)) mice. CD45.1/CD45.2*Scl-CreJak2^VF^* mice were generated by crossing Scl*-CreJak2^VF^* mice to CD45.1 (The Jackson Laboratory B6.SJL-Ptprca Pepcb/BoyJ (002014)) mice. CD45.1/CD45.2*Scl-Cre-Jak2^VF^*mice were crossed with ZsGreen reporter mice (The Jackson Laboratory, B6.Cg-Gt(ROSA)26Sor^tm6(CAG-ZsGreen1)Hze^/J (Ai6(RCL-ZsGreen) (007906) to generate CD45.1/CD45.2*Scl-CreJak2^VF^ZsGreen* mice. *Ldlr^−/-^*recipient mice for the *Mx1-CreJak2^VF^* moderate cholesterol lowering model that included *Myh11-Cre^ERT2^* and *ZsGreen* alleles from a previous lineage-tracing study were obtained from a collaborator and were previously described (33). However, these alleles were not relevant to the experimental outcomes and were not analyzed. Half of recipients were *Ldlr^−/-^ Myh-11-CreZsGreen* while the other half either lacked the *ZsGreen* allele or both the *ZsGreen* allele and the *Myh11-Cre*. For the *Mx1-CreJak2^VF^* and *Scl-CreJak2^VF^* aggressive cholesterol lowering studies, *Ldlr^−/-^* recipient mice were purchased from The Jackson Laboratory (B6.129S7-*Ldlr*tm1Her/J (002207)). Littermate control mice for studies involving *Mx1-CreJak2^VF^* mice contained either an allele with *Mx1-Cre* or the *Jak2^VF^* transgene, but not both. Littermate control mice for studies involving *Scl-CreJak2^VF^ZsGreen* mice contained alleles for *Scl-Cre* and *ZsGreen* but not the *Jak2^VF^*transgene.

### 2.3 Bone Marrow Transplantation

Bone marrow transplantations (BMT) were conducted as previously described with the following exceptions (10). For the *Mx1-CreJak2^VF^* moderate cholesterol lowering study, donor mice were 8-12 weeks old while recipients were 11-14 weeks old. For the *Mx1-CreJak2^VF^* effective cholesterol lowering study, donor mice were 7-16 weeks old while recipients were 7 weeks old. For the *Scl-CreJak2^VF^* study, donor mice were 6-12 weeks old while recipients were 8 weeks old. For the *Mx1-CreJak2^VF^* moderate cholesterol lowering study, recipients were lethally irradiated once with 10.5 Gy from a cesium gamma source. Due to decommissioning of the cesium gamma source, recipients from the *Mx1-CreJak2^VF^* and *Scl-CreJak2^VF^*effective cholesterol lowering studies were irradiated with 10.5 Gy using a Multirad 350 X-ray Irradiator from Precision X-Ray. Within 24 hours of irradiation, bone marrow was isolated from donors with the indicated genotypes. Total bone marrow cell number was quantified using an iN CYTO C-Chip hemocytometer (DHC-N01) according to the manufacturer instructions. Irradiated mice were randomized to treatment groups, anesthetized with isoflurane before receiving 3 × 10^6^ total bone marrow cells via intravenous (i.v.) injection at a total final volume of 100 µl. For the *Mx1-CreJak2^VF^*studies, recipients received either 6 × 10^5^ *CD45.1Mx1-CreJak2^VF/WT^* or 6 × 10^5^ *CD45.1* control bone marrow cells (20%) combined with 2.4 × 10^6^ *CD45.2Jak2^WT/WT^* cells (80%). For the *Scl-CreJak2^VF^* study, recipients received either 6 × 10^5^ *CD45.1CD45.2Scl-CreJak2^VF/WT^ZsGreen* or 6 × 10^5^ *CD45.1CD45.2Scl-CreJak2^WT/WT^ZsGreen* bone marrow cells (20%) combined with 2.4 × 10^6^ *CD45.1Jak2^WT/WT^* cells (80%). For the *Mx1-CreJak2^VF^*studies, mice were allowed to recover for four weeks after BMT before being injected intraperitonially (i.p.) with 50 µg/mouse/day polyinosinic:polycytidylic acid (pIpC) two times, 48 hours apart.

### 2.4 Atherosclerosis Studies

For the *Mx1-CreJak2^VF^* studies, power analysis using GPower3.1 indicated that with a total sample size of 80 (n = 20 per group), the study had 80% power to detect an effect size of f = 0.32 at α = 0.05, using two-way analysis of variance (ANOVA) across two genotypes and two treatments. For the *Scl-CreJak2^VF^* study, power analysis using GPower3.1 indicated that with a total sample size of 80 (n = 16 per group), the study had 80% power to detect an effect size of f = 0.40 at α = 0.05, using one-way ANOVA across five groups. Four weeks after BMT, *Ldlr^−/-^* recipient mice were fed a western diet (ENVIGO, cat. no. TD.88137) for the indicated times. For the final 10 days of the western diet in the *Scl-CreJak2^VF^* study, mice were switched to a western diet containing tamoxifen at 500 mg/kg (ENVIGO, cat. no. TD130889). To induce LDL lowering in the *Mx1-CreJak2^VF^* study of Figure 1A, mice were administered 2 × 10^11^ viral particles per mouse in 200 µl of 1X PBS via tail vein injection of an hDAD-*hLDLR* vector (Gene Vector Core Laboratory at the Baylor College of Medicine) and switched to chow diet. To induce LDL lowering in the *Mx1-CreJak2^VF^* study of Figure 4A and the *Scl-CreJak2^VF^* study of Figure 6A, mice were administered 2.5 × 10^11^ genome copies per mouse in 100 µl of 1X PBS via tail vein injection of an AAV8-TBG-*mLdlr* vector (AAV-263355, Vector Biolabs) and switched to chow diet. Following euthanasia, aortic roots were collected and fixed with 4% paraformaldehyde (PFA) (Electron Microscopy Sciences 15710; 16% PFA diluted 1:4 in 1.33X PBS) for 24 hours at 4^°^C, then embedded in paraffin.

**Figure 1.**
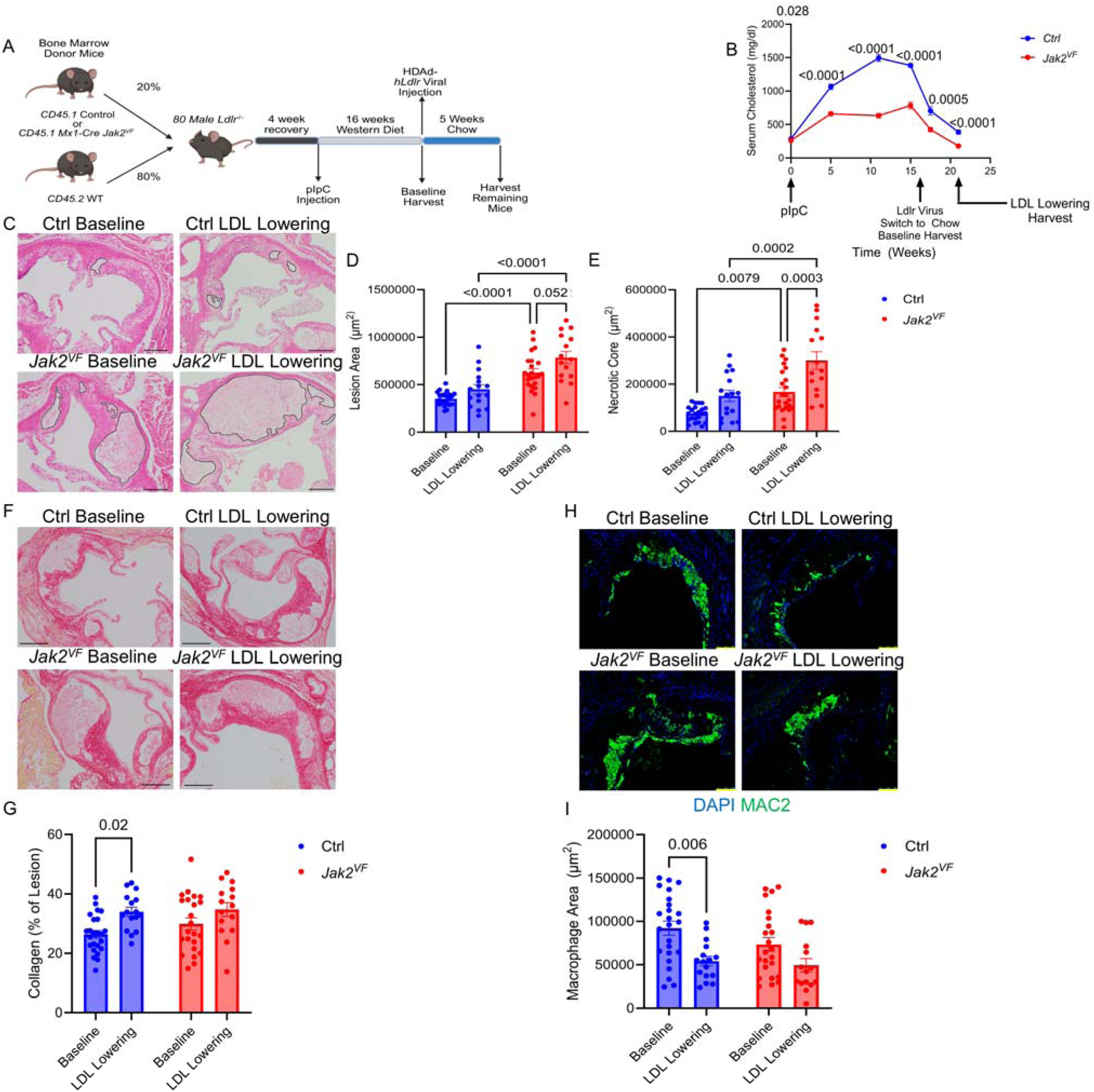
Impaired Atherosclerosis Regression in *Jak2^VF^*MPN Mice with Moderate Cholesterol Lowering. **A.** Study Design, created with BioRender.com. **B.** Plasma cholesterol (n = 34, 55, 55, 54, 16, 16 for Ctrl mice, n=31, 55, 49, 39, 17, 16 for *Jak2^VF^* mice, for weeks 0, 5, 11, 15, 17.5, 21 respectively). *P* = 0.028, <0.0001, <0.0001, <0.0001, 0.0005, <0.0001 (Ctrl vs *Jak2^VF^* at weeks 0, 5, 11, 15, 17.5, 21 respectively). **C.** H&E images of aortic root lesions. Black lines, necrotic core. Scale bar, 200 µm. **D.** Lesion area, n = 15-23. *P* < 0.0001 (Ctrl Baseline vs *Jak2^VF^* Baseline; Ctrl LDL Lowering vs. *Jak2^VF^*LDL Lowering), *P* = 0.052 (*Jak2^VF^* Baseline vs LDL Lowering). **E.** Necrotic core area, n = 15-23. *P* = 0.0079 (Ctrl Baseline vs *Jak2^VF^* Baseline), *P =* 0.0002 (Ctrl LDL Lowering vs. *Jak2^VF^* LDL Lowering), *P* = 0.0003 (*Jak2^VF^*Baseline vs LDL Lowering). **F.** Picrosirius red-stained aortic root lesions. Scale bar, 278 µm **G.** Collagen area as a percentage of lesion area, n = 15-24. *P* = 0.02 (Ctrl Baseline vs LDL Lowering). **H.** Images of aortic root lesions stained for MAC2 (Green) and DAPI (Blue). Scale bar, 100 µm. **I.** Macrophage area, n = 15-24. *P* = 0.006 (Ctrl Baseline vs LDL Lowering). All quantifications shown as mean ± s.e.m. Two-way ANOVA with the Geisser-Greenhouse correction for sphericity and Tukey’s multiple comparisons test (B). Two-way ANOVA with Tukey’s multiple comparisons test (D,E,G,I).

### 2.5 Histological Analysis

Lesion area and necrotic core area were quantified as previously described (22,30). Blinded researchers sectioned paraffin-embedded aortic root lesions at similar points between mice over a 150 µm span for a total of 50 sections spaced 3 µm apart with 2 sections per slide on 25 slides. Six total slides spaced 30 µm apart were then stained with hematoxylin & eosin (H&E) (Sigma MHS32-1L; Abcam ab246824) and imaged with an Olympus DP27 camera attached to an Olympus CX43 biological microscope set to the 10X objective using Olympus cellSens Entry 3.1 software. Lesion area was quantified in a blinded fashion, and the average of the six slides was used to determine lesion area. The Necrotic core area was similarly quantified in a blinded fashion using the same six H&E sections. Collagen was stained using the Polysciences Picrosirius Red Stain Kit (cat. no. 24901) according to the manufacturer’s instructions. The same slide between mice was stained and the collagen quantified by applying a consistent color threshold to calculate the collagen-positive area as a percentage of total lesion area. Picrosirius red was imaged via the same system used for H&E imaging. All lesion analysis was conducted using FIJI software.

### 2.6 Immunofluorescence

Identical slides of aortic roots between mice were baked in an incubator at 60^°^C for 30 minutes and deparaffinized in Histo-clear (National Diagnostics HS-200/50-899-90147) three times for 10 minutes. Histo-clear was removed by 20 shakes in 100% ethanol twice followed by 20 shakes in Histoprep Reagent Alcohol (fisher HC-600-1GAL) before being washed in tap water 7 times. Antigen retrieval was performed by placing slides either in citric acid-based antigen retrieval solution (Vector Laboratories, H3300) or tris-based antigen retrieval solution (Vector Laboratories, H3301) and pressure cooking for 15 minutes with a 5-minute natural release in an Instant Pot IP-LUX pressure cooker. Citric acid-based antigen retrieval was performed for all antibodies except for the antibody against cleaved GasD. After pressure cooking, slides were cooled for 10 minutes and then washed 3 times for 5 minutes in 1X phosphate buffered saline (PBS) (Boston Bioproducts BM-220; diluted 1:10). Hydrophobic circles were drawn using a liquid blocker pen (Sigma Z377821). For antibodies sourced from mice, mouse-on-mouse blocking was performed by incubating sections in 10% goat serum (ThermoFisher 10000C) in 1X PBS (Corning 21-040-CM) with 1 drop of mouse-on-mouse block per 1.25ml of solution (Vector Laboratories NC9290646) for 1 hour at room temperature in a humidified chamber. Slides were then washed in 1X PBS containing tween 20 (Fisher BP337-500) at a concentration of 0.01% (PBS-T) twice for 5 minutes and then in 1X PBS once for 5 minutes. For other antibodies, sections were blocked by incubating in Normal Serum Block (Biolegend 927503) for 1 hour at room temperature in a humidified chamber. Following blocking, sections were incubated with the following primary antibodies at the indicated concentrations overnight at 4^°^C in a humidified chamber: Absent in melanoma 2 (AIM2) (Abcam, ab119791, 1:250), Cleaved GasD (Cell Signaling, 10137, 10µg/ml), c-myc (Cell Signaling, 5605, 5.76 µg/ml), Ki67 (Abcam, ab15580, 9 µg/ml), MAC2 conjugated to Alexa Fluor 488 (Cedarlane, CL8942AF4, 1 µg/ml), MAC2 (Cedarlane, CL8942AP, 1 µg/ml), mer proto-oncogene tyrosine kinase (MerTK) (R&D, BAF591, 2 µg/ml), pγH2AX (Cell Signaling, 9718, 0.74 µg/ml), ZsGreen (ThermoFisher, TA180002, 10 µg/ml), triggering receptor expressed on myeloid cells 2 (TREM2) (Denali, 4D9 DC1847, 1:100). The following IgG antibodies were used as negative controls for primary antibodies at the same concentration: Mouse IgG (ChromPure, 015-000-003), Rabbit IgG (Abcam, ab172730), Rabbit IgG (Novus Biologicals, NBP2-24891), Rabbit IgG (ChromPure 011-000-003), Rat IgG (ChromPure, 012-000-003), Rat IgG (Biolegend, 400431), Rat IgG conjugated to Alexa Fluor 488 (Invitrogen, 53-4321-80). Slides were then washed twice with PBS-T for 5 minutes and once with 1X PBS for 5 minutes. The following secondary antibodies were used at a dilution of 1:200 for 1 hour at room temperature: Rat conjugated to Alexa Fluor 488 (Invitrogen A11006), Rat conjugated to Alexa Fluor 568 (Invitrogen A11077), Rat conjugated to Alexa Fluor 647 (Invitrogen A-21247), Rabbit conjugated to Alexa Fluor 568 (Invitrogen A11011), Rabbit conjugated to Alexa Fluor 647 (A27040). The following secondary antibodies were used at a dilution of 1:800 for 1 hour at room temperature: Mouse conjugated to Alexa Fluor 647 (Invitrogen A21237), Streptavidin conjugated to Alexa Fluor 568 (Invitrogen S11226), Streptavidin conjugated to Alexa Fluor 647 (Invitrogen S32357). DAPI (4’,6-Diamidino-2-Phenylindole, Dilactate) (Biolegend, 422801) was included during incubation with secondary antibodies at a concentration of 5 µg/ml. Slides were then washed twice with PBS-T for 5 minutes and once with 1X PBS for 5 minutes. After washing, slides were mounted using Prolong Gold Antifade Reagent with DAPI (Invitrogen P36935). TUNEL staining was performed using the In Situ Cell Death Detection Kit TMR Red (Roche, 12156792910) according to the manufacturer’s instructions. Slides were imaged on a Leica Fluorescent DMI 6000B wide-field microscope or a Nikon Ti Eclipse inverted microscope with high-sensitivity confocal imaging at the Confocal and Specialized Microscopy Shared Resource (CSMSR) Core at Columbia University. All images were analyzed in a blinded fashion using FIJI software.

### 2.7 Blood Isolation and Analysis

Blood was collected at the indicated timepoints via retro-orbital sinus using heparinized microhematocrit capillary tubes (Fisherbrand 22-362566) into EDTA-coated microvette tubes (Sarstedt 20.1278.100). Complete blood cell counts were quantified using a VetScan HM5 Hematology system (Zoetis/Abaxis). Plasma was collected by centrifuging blood at 13,000 x g for 10 minutes at 4^°^C. Plasma cholesterol was quantified using the Cholesterol E kit (Fujifilm Wako Pure Chemical Corporation, #999-02601/NC9138103) according to the manufacturer’s instructions. Flow cytometric analysis for blood leukocytes was performed by lysing in RBC lysis buffer (Biolegend 420302) for 2 minutes at room temperature before centrifuging at 3,000 x g for 2 minutes at 4^°^C. Cells were washed in ice cold MACS Buffer (0.5% BSA and 2mM EDTA in 1X PBS) (FischerScientific BP9706-100, Invitrogen 15575-038) and centrifuged at 800 x g for 10 minutes at 4^°^C before staining with antibodies at a 1:100 dilution in ice cold MACS buffer for 20 minutes. Cells were washed once more in ice cold MACS Buffer, centrifuged at 800 x g for 10 minutes at 4^°^C before being resuspended in 250 µl of MACS Buffer. Antibodies used in each experiment are listed in Supplementary Tables S1–S3. Cells were captured using a BD Fortessa Flow Cytometer running BD FACSDiva software at the Flow Cytometry Shared Core of the Columbia Center for Translational Immunology (CCTI) and Herbert Irving Comprehensive Cancer Center (HICCC). Flow cytometric data were analyzed using FCS Express 7 Research Edition.

### 2.8 Bone Marrow-Derived Macrophage Cultures

Hindlimbs were isolated and stored in ice cold DMEM. The ends of the bones were cut and the bones placed into 600 µl microcentrifuge tubes with a hole punctured at the bottom and sitting inside of 1.5 ml microcentrifuge tubes containing 100 µl of ice cold DMEM. The bones were pulse centrifuged at 4^°^C for 8 seconds. The pellet was then suspended in 1.5 ml of ice cold DMEM before being filtered through a 40 µm cell strainer (Fisher 22-363-547) into 50 ml of ice cold DMEM. Suspensions were centrifuged at 800 x g for 10 minutes at 4^°^C and the supernatant aspirated. The pellet was resuspended in M-CSF complete media for the *Jak2^VF^Aim2^−/-^* experiment (DMEM + 20 ng/ml M-CSF + 10% FBS + 1% PENSTREP) (M-CSF from Cell Signaling, 33444S or LCM complete media (all other experiments) (DMEM + 20% LCM + 10% FBS + 1% PENSTREP) and plated into tissue culture treated dishes (Corning CLS430167). Cells were differentiated into macrophages by incubating for 5 days at 37^°^C and 5% CO_2_. After 5 days, the media was aspirated and cells washed in 20 ml of 1X PBS before being stripped by incubation in 5 ml of Corning Cell Stripper (Corning 25-056-CL) for 5 minutes before being collected using 5 ml of complete media. Cells were then centrifuged at 800 x g for 10 minutes at 4^°^C and the supernatant aspirated. Cells were counted using an iN CYTO C-Chip hemocytometer (DHC-N01) according to the manufacturer instructions. Cells were plated into well plates of various sizes depending on experiment at a density of 333,333 cells/ml (3 ml of suspension for 6-well plates, 2 ml of suspension for 12-well plates). Cells were allowed to reattach overnight before proceeding with further experiments.

### 2.9 acLDL Cell Culture Experiments

Preparation of acetyl-LDL (acLDL) was performed as previously described (34). Bone marrow-derived macrophages (BMDMs) were differentiated and plated as described in section 2.8 and then incubated in M-CSF-free media containing either vehicle or 25 µg/ml acLDL overnight at 37^°^C and 5% CO_2_.

### 2.10 oxPAPC Cell Culture Experiments

BMDMs were differentiated and plated as described in section 2.8 and then incubated in M-CSF-free media containing either vehicle or 50 µg/ml oxidized 1-palmitoyl-2-arachidonoyl-sn-glycero-3-phosphocholine (oxPAPC) overnight at 37^°^C and 5% CO_2_.

### 2.11 M**β**CD-Cholesterol Cell Culture Experiments

BMDMs were differentiated and plated as described in 2.8. Cells were then primed in complete media containing 100 ng/ml lipopolysaccharide (LPS) (Cell Signaling 14011) for 3 hours at 37^°^C and 5% CO_2_ before treating with complete media containing vehicle, (2-(2,2,6,6-tetramethylpiperidin-1-oxyl-4-ylamino)-2-oxoethyl)triphenylphosphonium chloride (mitoTEMPO) (MedChemExpress HY-112879) (10 µM), methyl-β-cyclodextrin-cholesterol (MβCD-Cholesterol) (Sigma C4951) (30 µg/ml) or a combination of mitoTEMPO (10 µM) and MβCD -Cholesterol (30 µg/ml) for 4 hours at 37^°^C and 5% CO_2_. LPS remained in the media for the duration of these treatments at 100 ng/ml.

### 2.12 Bone Marrow-Derived Macrophage Proliferation Quantification

BMDM proliferation was measured using the Invitrogen Click-iT™ Plus EdU Alexa Fluor™ 488 Flow Cytometry Assay Kit according to the manufacturer’s instructions. After incubation with EdU, cells were washed in 1X PBS before being stripped by incubation in 500 µl of Corning Cell Stripper (Corning 25-056-CL) for 5 minutes. Cells were captured using a BD Fortessa Flow Cytometer running BD FACSDiva software at the Flow Cytometry Shared Core of the Columbia Center for Translational Immunology (CCTI) and Herbert Irving Comprehensive Cancer Center (HICCC). Flow cytometric data were analyzed using FCS Express 7 Research Edition.

### 2.13 Immunoblotting

Cells were lysed in radioimmunoprecipitation assay (RIPA) buffer containing protease and phosphatase inhibitors (Invitrogen 87786, ThermoFisher 78420) for 30 minutes at 4^°^C on a rocker. Lysates were then frozen at -80^°^C to enhance lysis. Lysates were then thawed at 4^°^C, centrifuged at max speed for 5 minutes at 4^°^C before being sonicated at 50% power for 30 seconds. Total protein concentration was determined by h bicinchoninic acid (BCA) analysis (ThermoFisher Scientific, 23225) according to the manufacturer’s instructions. Equal protein amounts were then loaded into polyacrylamide gels (Biorad 5671094) using 1X Running Buffer (Boston Bioproducts BP-150) and transferred to nitrocellulose membranes (Biorad 1620115) using transfer buffer (Boston Bioproducts BP-190) with 20% ethanol (Fisher 2701).

Membranes were washed in Tris-Buffered Saline with Tween 20 (TBS-T) (Boston Bioproducts IBB-180X) for 5 minutes at room temperature before blocking for 1 hour at room temperature in 3% bovine serum albumin (BSA) (Fisher BP9706-100) in TBS-T while gently rocking. After blocking, membranes were incubated overnight at 4^°^C in 1:1000 primary antibody solution in 3% BSA in TBS-T with the following antibodies: pYH2AX (Cell Signaling 9718) and total GasD (Cell Signaling 38754) or for 20 minutes at room temperature in 1:80,000 primary antibody solution with the following antibodies: β-Actin (Cell Signaling 12262). After incubation with non-HRP-conjugated primary antibodies, membranes were washed in TBS-T 5 times for 5 minutes before incubation in a horseradish peroxidase (HRP) conjugated anti-rabbit secondary antibody (7074 Cell Signaling Technology) in 3% BSA in TBS-T for 1 hour at room temperature while gently rocking. After HRP-conjugated primary or secondary antibody incubation, membranes were washed 5 times for 5 minutes before incubation for 30 seconds with a chemiluminescent substrate (ThermoFisher 34578). Membranes were then exposed in a dark room using film (Fisher XAR ALF 2025). Densitometric quantification was performed using FIJI software.

### 2.14 IL1**β** ELISA Detection

IL1β detection was quantified using the Mouse IL-1 beta/IL-1F2 DuoSet enzyme-linked immunosorbent assay (ELISA) (R&D Systems DY401) according to the manufacturer’s instructions. The plate was detected using a SpectraMax M2 (Molecular Devices).

### 2.15 Statistics

Statistical analyses were conducted using GraphPad Prism v10.5. Data are presented as the mean ± s.e.m. All statistical tests were performed using two-sided analyses. Normality was tested using the D’Agostino-Pearson omnibus and, where appropriate, log transformation was applied to approximate normality. Repeated measures data were analyzed using two-way analysis of variance (ANOVA) with the Geisser-Greenhouse correction for sphericity and Tukey’s multiple comparisons test with individual variance estimation. For datasets involving two independent variables, two-way ANOVA with Tukey’s multiple comparisons test or Kruskal-Wallis with Dunn’s multiple comparisons test were used based on distribution. For datasets involving five experimental groups, one-way ANOVA with Holm-Sidak’s multiple comparisons test or Kruskal-Wallis with Dunn’s multiple comparisons test were used based on distribution. Formal outliers were removed using a two-tailed Grubbs’ test (α = 0.05). For experiments with technical replicates from a single mouse, 3–6 wells were plated per condition. Wells were excluded as technical outliers when 6 wells were plated if the coefficient of variation (%CV) among wells for a given condition exceeded 20% (predefined QC threshold).

## 3. Results

### 3.1 Atherosclerosis Regression is Impaired with Moderate Cholesterol Lowering in *Jak2^VF^*Mice

To assess the effect of *Jak2^VF^* MPN on atherosclerosis regression, we placed *Jak2^VF^* MPN male mice and controls on a western diet for 16 weeks and then induced cholesterol lowering using an hDAD vector containing the human LDL receptor gene (hDAD-*hLDLR*) combined with a switch to chow diet for an additional 5 weeks (Figure 1A). Prior studies were performed in female mice (16,30). *Jak2^VF^* MPN male mice exhibited splenomegaly, lower body weight, and increased RBCs, hematocrit, red blood cell distribution width (RDWc), and neutrophils, mirroring previous findings in females (Supplemental Figure 1A-I) (30). In this model, CD45.1+ cells are *Jak2^VF^* in experimental mice and WT in controls. CD45.1+ neutrophils and monocytes were elevated while CD45.1+ lymphocytes were reduced (Supplemental Figure 1J-L). Plasma cholesterol was reduced in *Jak2^VF^* MPN mice on Western diet (Figure 1B) similar to previous findings in female mice and humans (24,30,35). Plasma cholesterol remained moderately elevated for 10 days post-virus treatment but returned to pre-diet levels (∼200-300 mg/dl) by the end of the LDL lowering period (Figure 1B). At baseline, lesion area and necrotic core area were significantly increased in the aortic root of male *Jak2^VF^* MPN mice compared to controls, confirming that *Jak2^VF^* MPN worsens atherosclerosis in both male and female mice (Figure 1C-E) (30). Moderate cholesterol lowering halted progression of lesion and necrotic core area in control mice whereas *Jak2^VF^* MPN mice developed significantly larger necrotic cores and displayed a trend toward increased lesion area (p=0.052) (Figure 1C-E). Consistent with previous studies (36,37), lesional collagen was increased in control mice after cholesterol lowering with no significant change in *Jak2^VF^* MPN mice, further indicating impaired resolution in the *Jak2^VF^* MPN mice (Figure 1F,G). The most consistent phenotype found in atherosclerosis regression studies has been a decrease in macrophage burden (13,17,18,19,37). Macrophages were significantly decreased in control mice but not in *Jak2^VF^* MPN mice after cholesterol lowering (Figure 1H,I). These results indicate that atherosclerosis resolution is impaired in *Jak2^VF^* MPN with moderate cholesterol lowering.

We next assessed inflammatory features in plaques. Necrotic, inflammatory cell death is mediated by activated Gasdermin D (GasD) downstream of inflammasome activation (38). We used an antibody to the active N-terminal fragment of GasD to assess pyroptosis in plaques (29). Total cleaved GasD in plaques as well as cleaved GasD in the necrotic core were increased in *Jak2^VF^* MPN lesions at baseline and were not reduced by cholesterol lowering (Figure 2A-C). However, macrophage-specific cleaved GasD was decreased by cholesterol lowering in *Jak2^VF^* MPN lesions, suggesting that macrophage pyroptosis had been suppressed by the time of harvest (Figure 2D,E). Consistent with earlier findings in females, absent in melanoma 2 (AIM2)+ macrophages and phosphorylated histone H2A.X (pγH2AX), an indicator of double stranded DNA break formation, were increased in *Jak2^VF^* MPN lesions at baseline and were decreased by cholesterol lowering, suggesting reversal of AIM2 inflammasome activation (Figure 2F-I) (30). Additionally, macrophage proliferation trended higher at baseline in *Jak2^VF^* MPN mice and was significantly reduced by cholesterol lowering (Figure 2J,K). These data suggest reversal of macrophage AIM2 inflammasome activation and pyroptosis by moderate cholesterol lowering in *Jak2^VF^* MPN mice, correlating with decreased macrophage proliferation and decreased DNA damage response. The persistence of cleaved GasD in the necrotic core area may reflect ongoing pyroptosis during the early phase of LDL lowering when cholesterol levels were persistently elevated.

**Figure 2.**
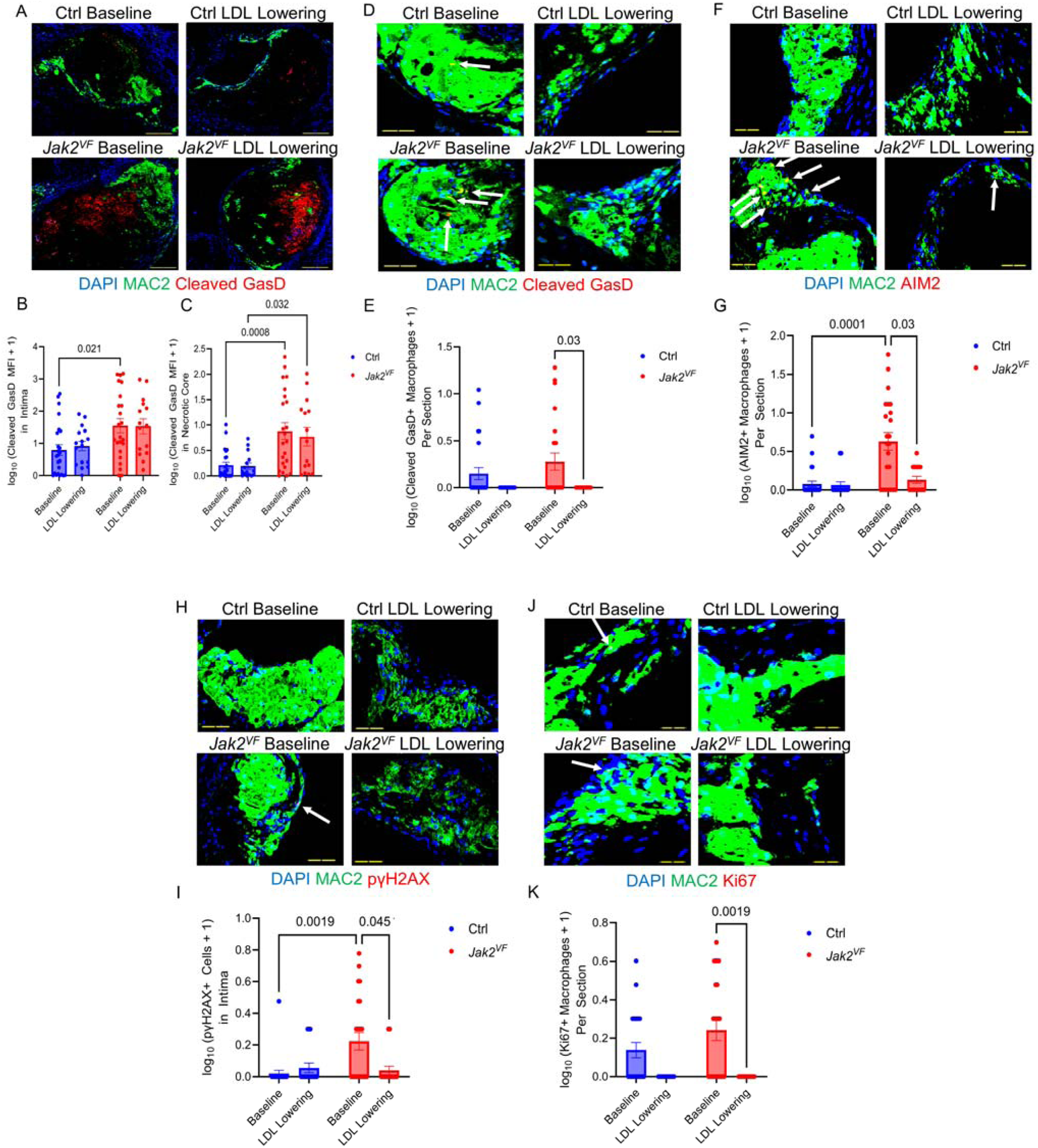
Moderate Cholesterol Lowering Reverses Macrophage AIM2 Inflammasome Activation, DNA Damage, and Proliferation in *Jak2^VF^*MPN Lesions. **A.** Images of aortic root lesions stained for MAC2 (Green), Cleaved GasD (Red), and DAPI (Blue). Scale bar, 127 µm. **B.** Log_10_ transformed cleaved GasD mean fluorescence intensity (MFI) in lesions with the addition of constant 1, n = 15-24. *P =* 0.021 (Ctrl Baseline vs *Jak2^VF^* Baseline). **C.** Log_10_ transformed cleaved GasD mean fluorescence intensity (MFI) in necrotic cores with the addition of the constant 1, n = 15-24. *P =* 0.0008 (Ctrl Baseline vs *Jak2^VF^* Baseline), *P =* 0.032 (Ctrl LDL Lowering vs *Jak2^VF^* LDL Lowering). **D.** Images of aortic root lesions stained for MAC2 (Green), Cleaved GasD (Red), and DAPI (Blue). Scale bar, 40 µm. White arrows, cleaved GasD+ macrophages. **E.** Log_10_ transformed cleaved GasD positive macrophages per section with the addition of constant 1, n = 15-24. *P =* 0.03 (*Jak2^VF^* Baseline vs LDL Lowering). **F.** Images of aortic root lesions stained for MAC2 (Green), AIM2 (Red), and DAPI (Blue). Scale bar, 75 µm. White arrows, AIM2 positive macrophages. **G.** Log_10_ transformed AIM2 positive macrophages per section with the addition of constant 1, n = 15-23. *P =* 0.0001 (Ctrl Baseline vs *Jak2^VF^* Baseline), *P =* 0.03 (*Jak2^VF^* Baseline vs LDL Lowering). **H.** Images of aortic root lesions stained for MAC2 (Green), pγH2AX (Red), and DAPI (Blue). Scale bar, 46 µm. White arrows, pγH2AX positive cells. **I.** Log_10_ transformed pγH2AX positive macrophages per section with the addition of constant 1, n = 15-23. *P =* 0.0019 (Ctrl Baseline vs *Jak2^VF^* Baseline), *P =* 0.045 (*Jak2^VF^* Baseline vs LDL Lowering). **J.** Images of aortic root lesions stained for MAC2 (Green), Ki67 (Red), and DAPI (Blue). Scale bar, 25 µm. White arrows, Ki67 positive macrophages. **K.** Log_10_ transformed Ki67 positive macrophages per section with the addition of constant 1, n = 15-23. *P =* 0.0019 (*Jak2^VF^* Baseline vs LDL Lowering). **L.** Images of aortic root lesions stained for MAC2 (Green), 8OHdG (Red), and DAPI (Blue). Scale bar, 78 µm. **M.** Percentage of MAC2 positive area double positive for 8OHdG and MAC2, n = 14-24. *P =* 0.023 (Ctrl Baseline vs LDL Lowering). MAC2 intensity differences reflect independent staining and imaging sessions across panels. All quantifications shown as mean ± s.e.m. Two-way ANOVA with Tukey’s multiple comparisons test (B,C). Kruskal-Wallis test with Dunn’s multiple comparisons test (E,G,I,K,M).

### 3.2 Cholesterol Lowering Restores Impaired Efferocytosis and increases TREM2+ Macrophages in *Jak2^VF^* mice

LDL Lowering has been associated with improvements in pro-resolving features of lesions such as increased macrophage efferocytosis, mer proto-oncogene tyrosine kinase (MerTK) expression (13,39,40), increased numbers of triggering receptor expressed on myeloid cells 2 (TREM2)+ macrophages and limitation of necrotic core formation in plaques (16,41). During atherosclerosis progression, macrophage MerTK and TREM2 were suppressed in *Jak2^VF^* CH mouse plaques as a result of interleukin-1 (IL-1) signaling from mutant to wild-type (WT) cells (29). Consistent with these studies, we found that macrophage MerTK was decreased in *Jak2^VF^*MPN lesions at baseline compared to controls (Figure 3A,B). MerTK levels were completely restored by cholesterol lowering (Figure 3A,B). Additionally, we found that cholesterol lowering dramatically increased the percentage of macrophages positive for TREM2 in both control and *Jak2^VF^*MPN lesions (Figure 3C,D). Consistent with these findings, macrophage-associated TUNEL+ nuclei as a percentage of total lesional TUNEL+ nuclei, an estimate of in situ efferocytosis (14), were significantly decreased in *Jak2^VF^* MPN lesions at baseline and restored by cholesterol lowering, (Figure 3E,F). Together, these data suggest that cholesterol lowering restores impaired efferocytosis in *Jak2^VF^* MPN lesions and increases the proportion of inflammation-resolving macrophages in lesions. Thus, macrophage inflammasome activation and associated defects in efferocytosis were reversed by cholesterol lowering in *Jak2^VF^* MPN mice. The increased necrotic core and decreased collagen content in these mice could reflect limited resolution of hypercholesterolemia during the earlier phases of LDL lowering.

**Figure 3.**
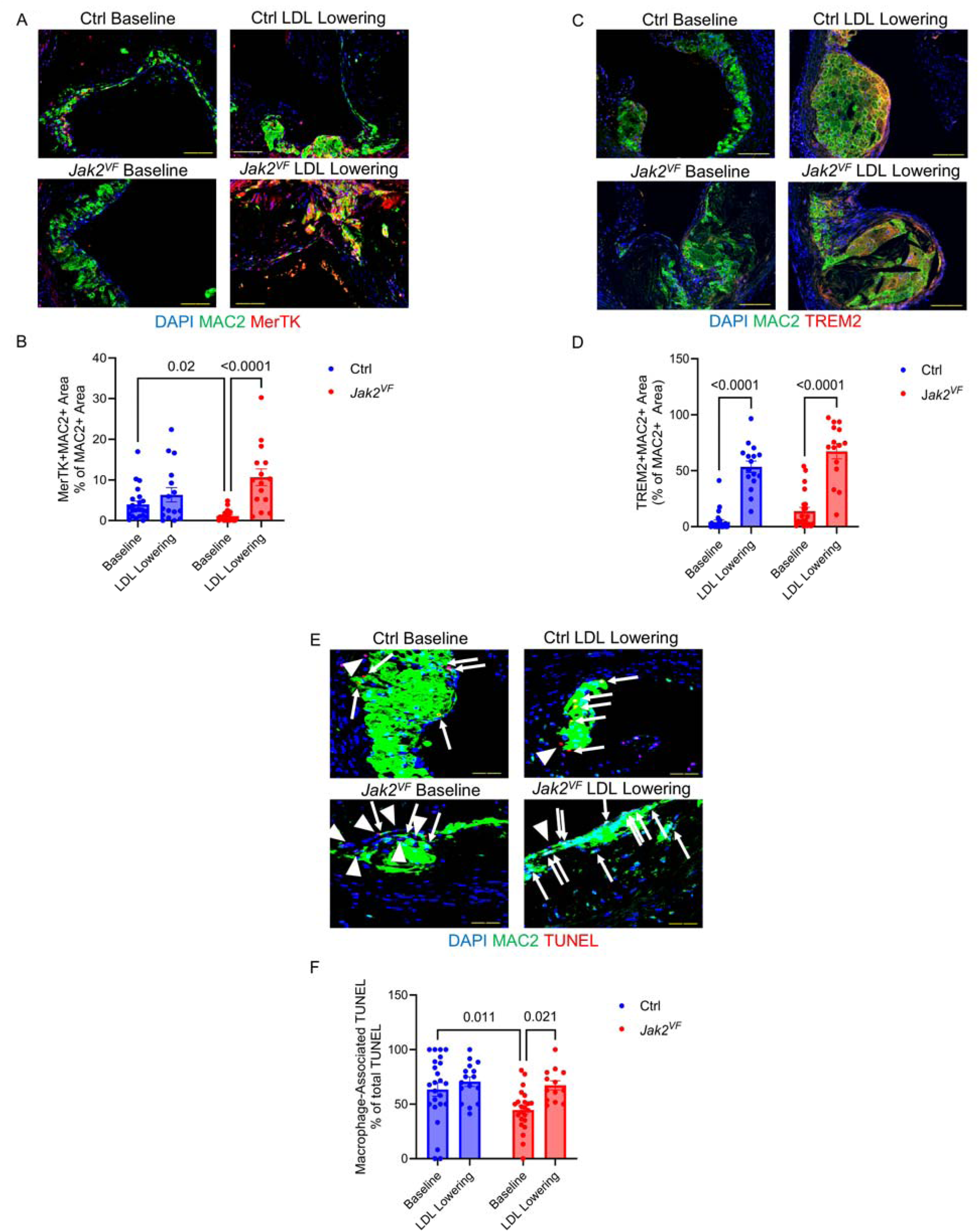
Moderate Cholesterol Lowering Reverses Impaired Efferocytosis in *Jak2^VF^*MPN Lesions While Increasing TREM2^Hi^ Macrophages in Control and *Jak2^VF^*MPN Lesions. **A.** Images of aortic root lesions stained for MAC2 (Green), MerTK (Red), and DAPI (Blue). Scale bar, 102 µm. **B.** Percentage of MAC2 positive area double positive for MerTK and MAC2, n = 15-23. *P =* 0.02 (Ctrl Baseline vs *Jak2^VF^*Baseline), *P* < 0.0001 (*Jak2^VF^* Baseline vs LDL Lowering). **C.** Images of aortic root lesions stained for MAC2 (Green), TREM2 (Red), and DAPI (Blue). Scale bar, 100 µm. **D.** Percentage of MAC2 positive area double positive for TREM2 and MAC2, n = 15-24. *P* < 0.0001 (Ctrl and *Jak2^VF^* Baseline vs LDL Lowering). **E.** Images of in situ efferocytosis in aortic root lesions: MAC2 (Green), TUNEL (Red), and DAPI (Blue). Scale bar, 40 µm. White arrows, nuclei double positive for TUNEL and MAC2. White wedges, nuclei positive for TUNEL but negative for MAC2. **F.** Percentage of TUNEL positive nuclei also positive for MAC2. *P =* 0.011 (Ctrl Baseline vs *Jak2^VF^* Baseline), *P* = 0.021 (*Jak2^VF^* Baseline vs LDL Lowering). MAC2 intensity differences reflect independent staining and imaging sessions across panels, n = 13-24. All quantifications shown as mean ± s.e.m. Two-way ANOVA with Tukey’s multiple comparisons test (D). Kruskal-Wallis test with Dunn’s multiple comparisons test (B,F).

### 3.3 More Effective Cholesterol Lowering Restores Atherosclerosis Regression

To test this hypothesis, we performed a similar study but with more effective LDL cholesterol lowering. Murine LDL does not bind as strongly to the human LDLR (employed in Figure 1) as to the mouse LDLR (42,43). Therefore, we employed an AAV8 containing the mouse *Ldlr* gene which more effectively and more quickly lowered cholesterol levels (Figure 4A). Blood parameters were similar to the first experiment and not appreciably changed by cholesterol lowering (Supplemental Figure 2, Figure 4B). Consistent with the first study, lesion area and necrotic core area were significantly increased in *Jak2^VF^* MPN mice at baseline. However, in contrast to the first study, while both groups showed non-significant upward trends of 1.29-fold (Ctrl) and 1.25-fold (*Jak2^VF^*) in lesion area, there was no change of necrotic core area in either group after aggressive LDL lowering (Figure 4C-E). Furthermore, aggressive cholesterol lowering significantly increased lesional collagen and decreased macrophage burden in both groups (Figure 4F-I). Analysis of cleaved GasD showed an increase in macrophages in *Jak2^VF^*mice at baseline that was reversed by cholesterol lowering, and in contrast to the first study a decrease in cleaved GasD in the necrotic core that appeared similar in control and *Jak2^VF^*MPN mice (Supplemental Figure 3A-D). Similarly, cleaved GasD was increased in CD11b+ splenocytes of *Jak2^VF^* MPN mice at baseline and reversed by cholesterol lowering (Supplemental Figure 3E,F). Consistent with the first study, double stranded DNA breaks (pγH2AX) were increased in *Jak2^VF^* MPN lesions at baseline and more strongly decreased by aggressive cholesterol lowering along with a significant decrease in macrophage proliferation (Supplemental Figure 3G-J). Additionally, macrophage MerTK was increased by aggressive cholesterol lowering (Supplemental Figure 3K,L). These results indicate that more effective cholesterol lowering completely reversed inflammatory features and induced similar levels of atherosclerosis resolution in control and *Jak2^VF^* MPN lesions.

### 3.4 Cholesterol Lowering Suppresses Proliferation of *Jak2^VF^* Macrophages and increasesTREM2+ Macrophages

In the previous studies *Jak2^VF^* was active during progression and lesion area was increased at baseline. To simulate the situation where the *Jak2^VF^* mutation was acquired later in life after atherosclerosis had already developed, we performed a third study in which *Jak2^VF^* was activated in HSCs at the time of LDL lowering by using a tamoxifen-inducible Scl-driven cre recombinase along with a floxed ZsGreen reporter allele (*Jak2^VF^ Scl* CH) to assess cell genotype specific changes (Figure 5A). In addition to the two LDL lowering groups, we included control and *Jak2^VF^* mice that remained on the western diet (Progression groups). As expected, plasma cholesterol remained elevated in the Progression groups but was markedly and rapidly decreased in both LDL lowering groups (Figure 5B). The *Jak2^VF^* MPN phenotype was partially apparent as indicated by increased spleen weights, RDWc, and hematocrit as well as expansion of *Jak2^VF^* (CD45.1+CD45.2+) blood monocytes and neutrophils, but less pronounced than in the earlier studies when *Jak2^VF^* was activated prior to Western diet feeding (Supplemental Figure 4).

**Figure 4.**
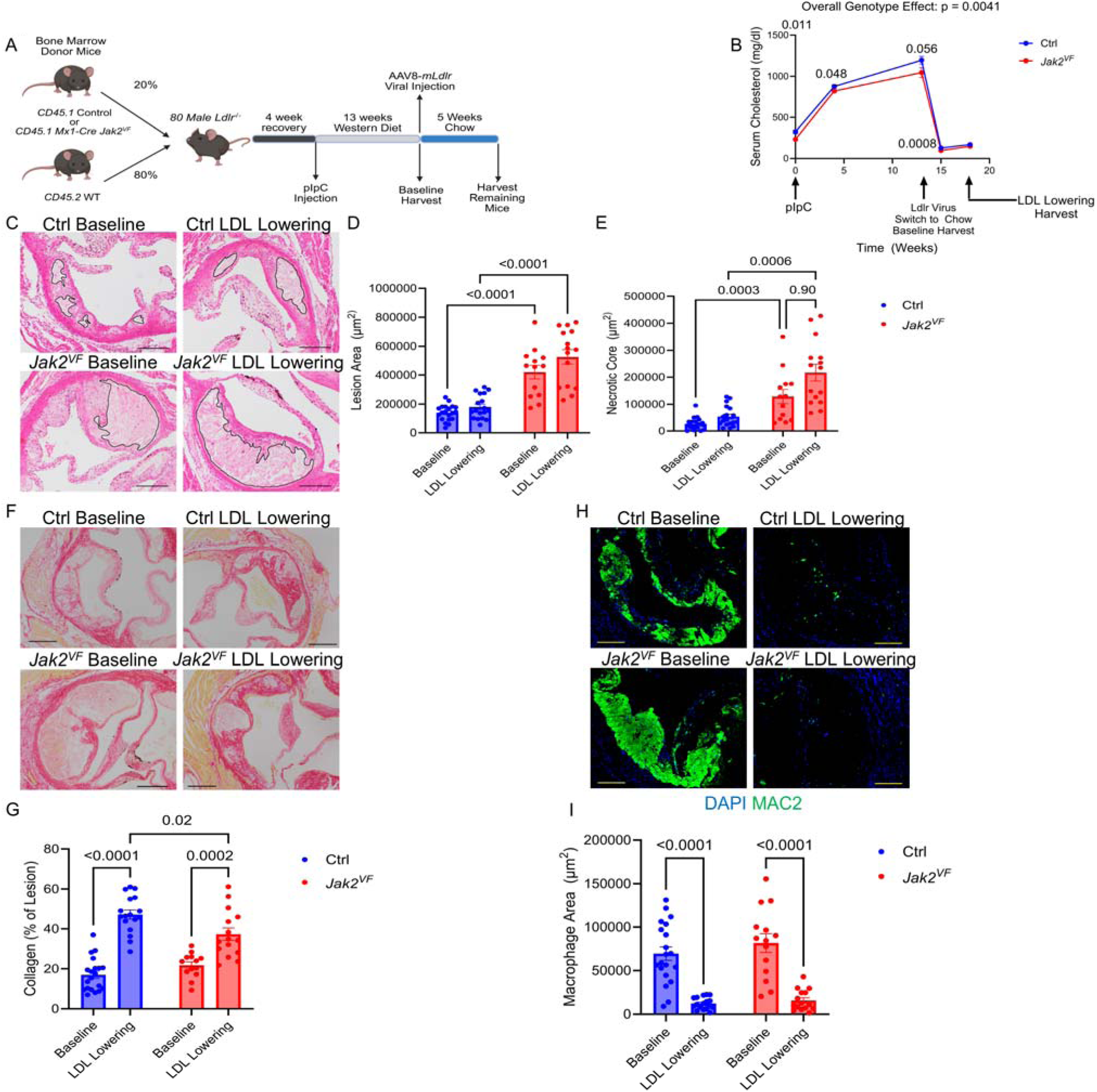
Aggressive Cholesterol Lowering Normalizes Regression in *Jak2^VF^* MPN Lesions. **A.** Study Design, created with BioRender.com. **B.** Plasma cholesterol (n = 5, 38, 19, 18, 18 for Ctrl mice, n = 5, 37, 15, 16, 15 for *Jak2^VF^* mice, for weeks 0, 4, 13, 15, 18 respectively). *P =* 0.0041 for genotype effect by two-way ANOVA with Geisser-Greenhouse correction, *P* = 0.011, 0.048, 0.056, 0.0008 (Ctrl vs *Jak2^VF^* at weeks 0, 4, 13, 15 respectively). **C.** H&E images of aortic root lesions. Black lines, necrotic core. Scale bar, 200 µm. **D.** Lesion area, n = 13-20. *P* < 0.0001 (Ctrl Baseline vs *Jak2^VF^* Baseline; Ctrl LDL Lowering vs. *Jak2^VF^* LDL Lowering). **E.** Necrotic core area, n = 13-20. *P* = 0.0003 (Ctrl Baseline vs *Jak2^VF^*Baseline), *P* = 0.0006 (Ctrl LDL Lowering vs. *Jak2^VF^* LDL Lowering), *P* = 0.90 (*Jak2^VF^* Baseline vs LDL Lowering). **F.** Picrosirius red-stained aortic root lesions. Scale bar, 219 µm **G.** Collagen area as a percentage of lesion area, n = 13-20. *P* < 0.0001 (Ctrl Baseline vs LDL Lowering), *P* = 0.0002 (*Jak2^VF^*Baseline vs LDL Lowering), *P* = 0.02 (Ctrl LDL Lowering vs. *Jak2^VF^* LDL Lowering). **H.** Images of aortic root lesions stained for MAC2 (Green) and DAPI (Blue). Scale bar, 193 µm. **I.** Macrophage area, n = 13-20. *P* < 0.0001 (Ctrl and *Jak2^VF^* Baseline vs LDL Lowering). All quantifications shown as mean ± s.e.m. Two-way ANOVA with Tukey’s multiple comparisons test (D,G,I). Two-way ANOVA with the Geisser-Greenhouse correction for sphericity and Tukey’s multiple comparisons test (B). Kruskal-Wallis test with Dunn’s multiple comparisons test (E).

**Figure 5.**
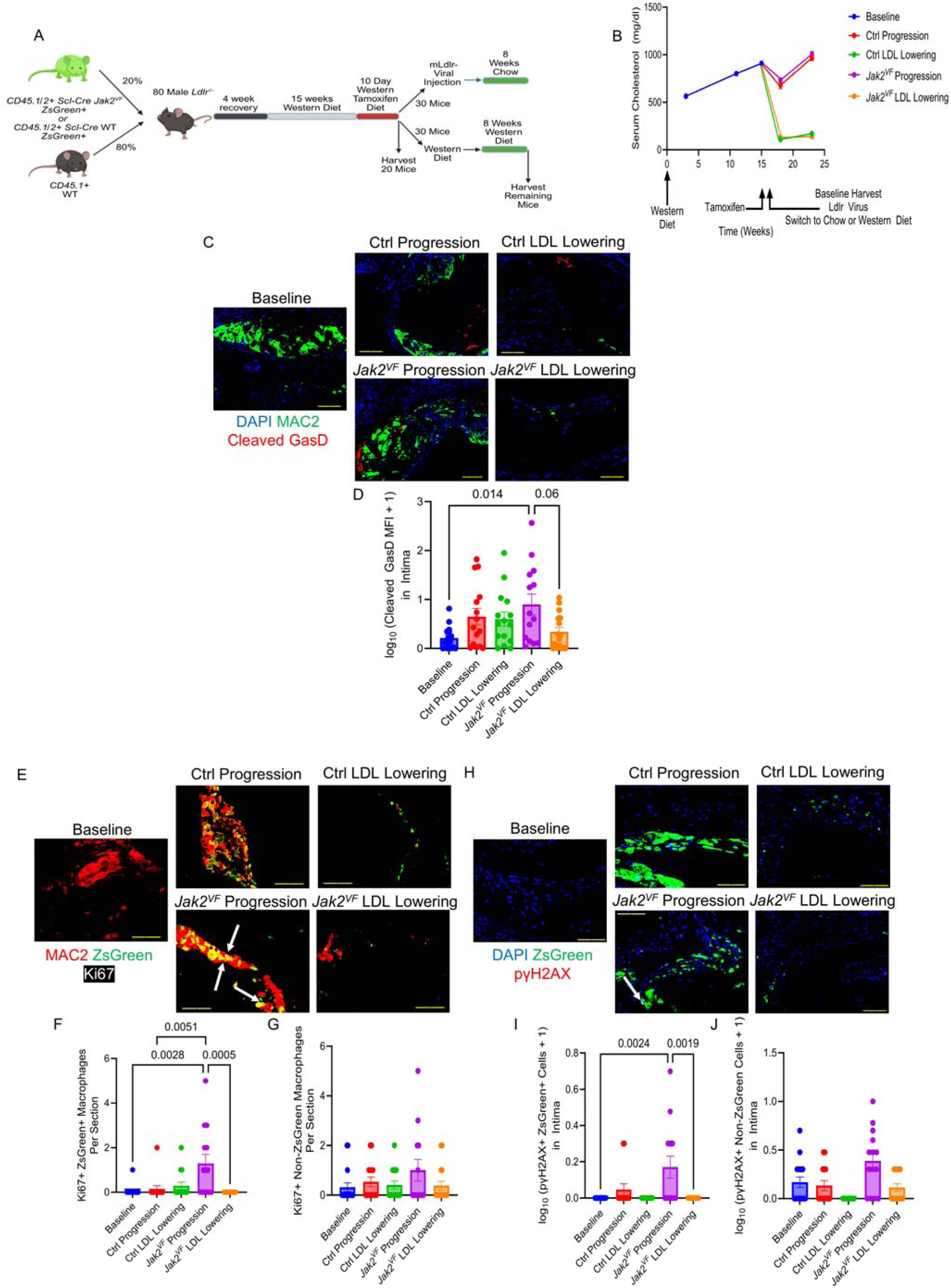
Cholesterol Lowering Suppresses *Jak2^VF^* Macrophage Proliferation and DNA Damage. **A.** Study design, created with BioRender.com. **B.** Plasma cholesterol (n = 80, 79, 79 for Baseline mice for weeks 3, 11, 15 respectively; n = 13, 15 for Ctrl Progression mice, n = 16, 16 for Ctrl LDL Lowering mice, n = 15, 15 for *Jak2^VF^* Progression mice, n = 16, 16 for *Jak2^VF^*LDL Lowering mice, for weeks 18, 23 respectively) **C.** Images of aortic root lesions for MAC2 (Green), Cleaved GasD (Red), and DAPI (Blue). Scale bar, 98 µm. **D.** Log_10_ transformed cleaved GasD mean fluorescence intensity (MFI) in lesions with the addition of constant 1, n = 14-16. *P* = 0.014 (Baseline vs *Jak2^VF^* Progression), *P* = 0.06 (*Jak2^VF^* Progression vs *Jak2^VF^* LDL Lowering). **E.** Images of aortic root lesions for MAC2 (Red), Ki67 (White), and ZsGreen (Green). Scale bar, 133 µm. White arrows, macrophages double positive for Ki67 and ZsGreen. **F.** Macrophages positive for both Ki67 and ZsGreen per section, n = 14-15. *P* = 0.0028 (Baseline vs *Jak2^VF^*Progression), *P* = 0.0051 (Ctrl Progression vs *Jak2^VF^* Progression), *P* = 0.0005 (*Jak2^VF^* Progression vs *Jak2^VF^*LDL Lowering). **G.** Macrophages positive for Ki67 but negative for ZsGreen per section, n = 13-16. **H.** Images of aortic root lesions stained for pγH2AX (Red), ZsGreen (Green), and DAPI (Blue). Scale bar, 75 µm. White arrows, pγH2AX positive cells. **I.** Log_10_ transformed cells double positive for pγH2AX and ZsGreen in lesions with the addition of constant 1, n = 13-16. *P* = 0.0024 (Baseline vs *Jak2^VF^* Progression), *P* = 0.0019 (*Jak2^VF^*Progression vs *Jak2^VF^* LDL Lowering). **J.** Log_10_ transformed cells positive for pγH2AX but negative for ZsGreen in lesions with the addition of constant 1, n = 13-16. All quantifications shown as mean ± s.e.m. One-way ANOVA with Holm-Sidak’s multiple comparisons test (D). Kruskal-Wallis test with Dunn’s multiple comparison’s test (F,G,I,J).

Compared to the earlier studies, the impact of *Jak2^VF^*on lesion progression was less pronounced reflecting the shorter period of *Jak2^VF^* activation (Supplemental Figure 5A-C). However, consistent with the earlier studies, the necrotic core significantly worsened in the *Jak2^VF^* high LDL mice but not in the control mice compared to baseline, while the necrotic core was not significantly changed in either LDL lowering group compared to baseline (Supplemental Figure 5C). Additionally, macrophage burden was similarly decreased in both LDL lowering groups, (Supplemental Figure 5D,E). These studies further support that aggressive cholesterol lowering similarly promotes resolution in control and *Jak2^VF^* mice with acquisition of *Jak2^VF^* in the context of established atherosclerosis.

Cleaved GasD was increased in *Jak2^VF^* lesions compared to baseline and trended lower (p=.06) after cholesterol lowering, consistent with aggressive cholesterol lowering reversing pyroptosis in lesions (Figure 5C,D). In the *Jak2^VF^* Progression group, proliferation was significantly increased in *Jak2^VF^* (green) macrophages but not in WT (non-green) macrophages indicating a mutant cell-specific effect, as seen previously (30). Importantly, this increased proliferation of *Jak2^VF^* cells was reversed by cholesterol lowering, (Figure 5E-G). The DNA damage response (pγH2AX) was significantly increased in *Jak2^VF^* cells and was reversed by cholesterol lowering while WT cells showed no significant proliferative changes or changes in the DNA damage response within *Jak2^VF^* lesions (Figure 5H-J).

Given that a proliferative efferocytic population with high c-Myc expression has been described with LDL lowering (13,39), we assessed c-Myc expression along with TREM2. In contrast to the decreases in Ki67, macrophage c-Myc and c-Myc+TREM2^Hi^ macrophages were increased by aggressive LDL lowering in both control and *Jak2^VF^* mice, suggesting a selective increase of TREM2+, anti-inflammatory macrophages despite a decrease in total macrophages (Figure 6).

### 3.5 Cholesterol Loading Increases AIM2 Inflammasome Activation and the DNA Damage Response in Macrophages

The parallel decrease in proliferation and DNA damage response in *Jak2^VF^* macrophages (Fig 5H-J) suggested that reversal of proliferation could be a key mechanism to explain reduced inflammasome activation in response to cholesterol lowering. To further assess the relationship between proliferation, the DNA damage response and inflammasome activation, we carried out mechanistic studies in bone marrow-derived macrophages (BMDMs). We found that cholesterol loading of macrophages with acetylated LDL (acLDL), which promotes entry of cholesterol via the scavenger receptor A/endo-lysosomal system (44), caused a modest increase in proliferation (Figure 7A). While *Jak2^VF^*macrophages showed significantly increased pγH2AX compared to controls, there was no effect of acLDL loading on pγH2AX (Figure 7B,C). We also showed that oxidized 1-palmitoyl-2-arachidonoyl-sn-glycero-3-phosphocholine (oxPAPC), a major component of oxidized LDL, stimulated macrophage proliferation, but this was associated with a reduced level of pγH2AX (Figure 7D-F). We next employed methyl-β-cyclodextrin (MβCD)-cholesterol, a distinctive mode of cholesterol loading that increases mitochondrial cholesterol content (45), promoting mitochondrial reactive oxygen species (ROS) formation and AIM2 inflammasome activation (46). MβCD-cholesterol increased IL-1β secretion into cell culture media especially in *Jak2^VF^* macrophages, and this was reversed by the mitochondrial-specific antioxidant mitochondria-targeted TEMPO (a triphenylphosphonium-conjugated nitroxide) (MitoTEMPO) (Figure 7G), suggesting that the effect was dependent on mitochondrial ROS production, consistent with prior studies (46). The increase in IL-1β secretion in *Jak2^VF^* macrophages in response to MβCD-cholesterol was reversed by AIM2 deficiency implicating the AIM2 inflammasome (Figure 7H). MβCD-cholesterol increased pγH2AX in control and *Jak2^VF^* macrophages, with a larger effect in the latter (two-way ANOVA) (Figure 7I,J). In contrast, MβCD-cholesterol loading did not increase macrophage proliferation (Figure 7K). Together these findings suggest that increased AIM2 inflammasome activation in response to MβCD-cholesterol loading may largely reflect increased mitochondrial ROS and DNA damage, rather than changes in cell proliferation.

**Figure 6.**
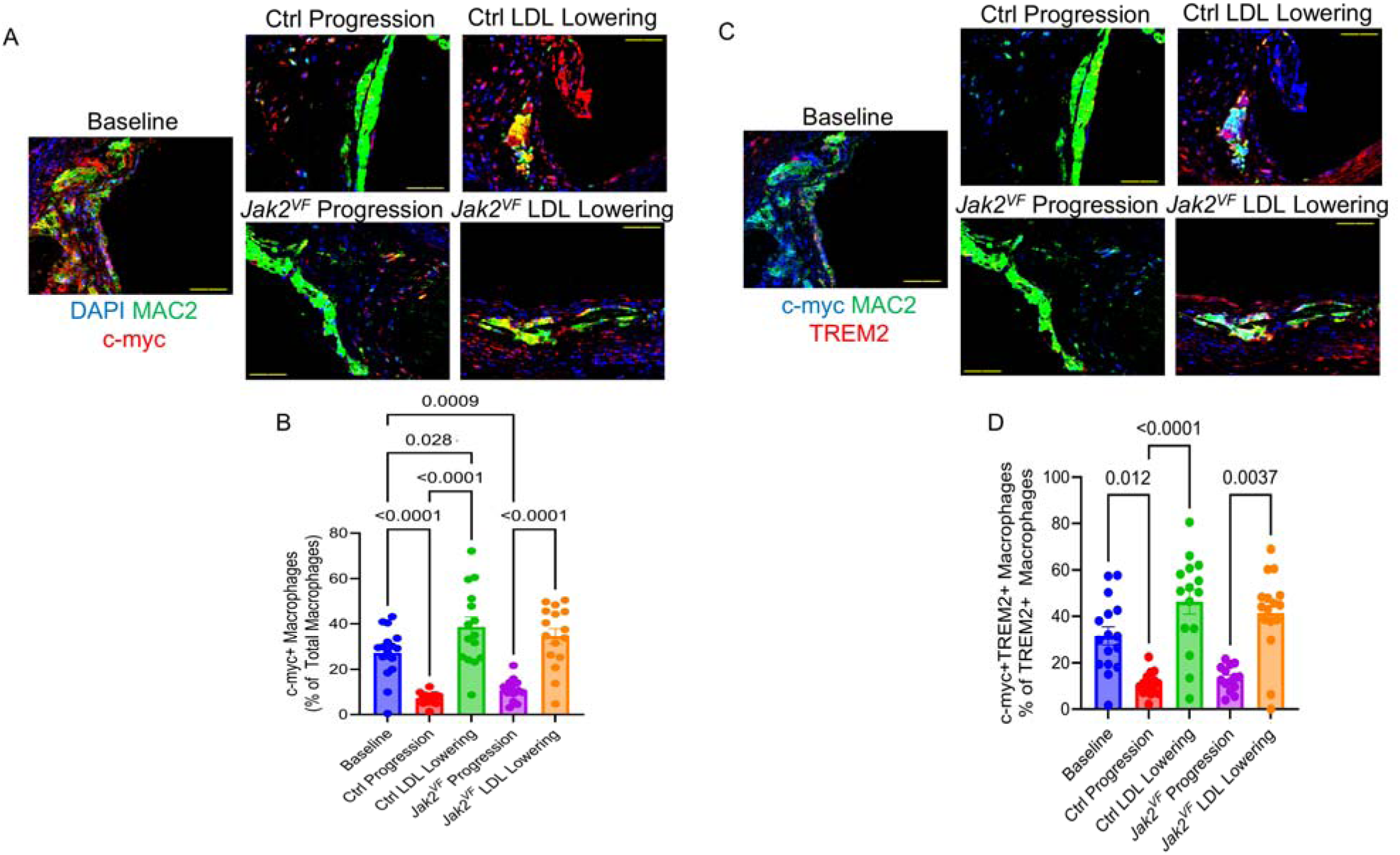
Cholesterol Lowering Increases c-myc in TREM2^Hi^ Macrophages. **A.** Images of aortic root lesions stained for MAC2 (Green), c-myc (Red), and DAPI (Blue). Scale bar, 121 µm. **B.** Percentage of macrophages positive for c-myc, n = 14-16. *P* < 0.0001 (Baseline vs Ctrl Progression; Ctrl Progression vs LDL Lowering; *Jak2^VF^* Progression vs LDL Lowering), *P* = 0.028 (Baseline vs Ctrl LDL Lowering), *P* = 0.0009 (Baseline vs *Jak2^VF^* Progression). **C.** Images of aortic root lesions stained for MAC2 (Green), c-myc (Blue), and TREM2 (Red). Scale bar, 121 µm. **D.** Percentage of TREM2 positive macrophages positive for c-myc, n = 14-16. *P* = 0.012 (Baseline vs Ctrl Progression), *P* < 0.0001 (Ctrl Progression vs LDL Lowering), *P* = 0.0037 (*Jak2^VF^* Progression vs LDL Lowering). All quantifications shown as mean ± s.e.m. One-way ANOVA with Holm-Sidak’s multiple comparisons test (B). Kruskal-Wallis test with Dunn’s multiple comparisons test (D).

**Figure 7.**
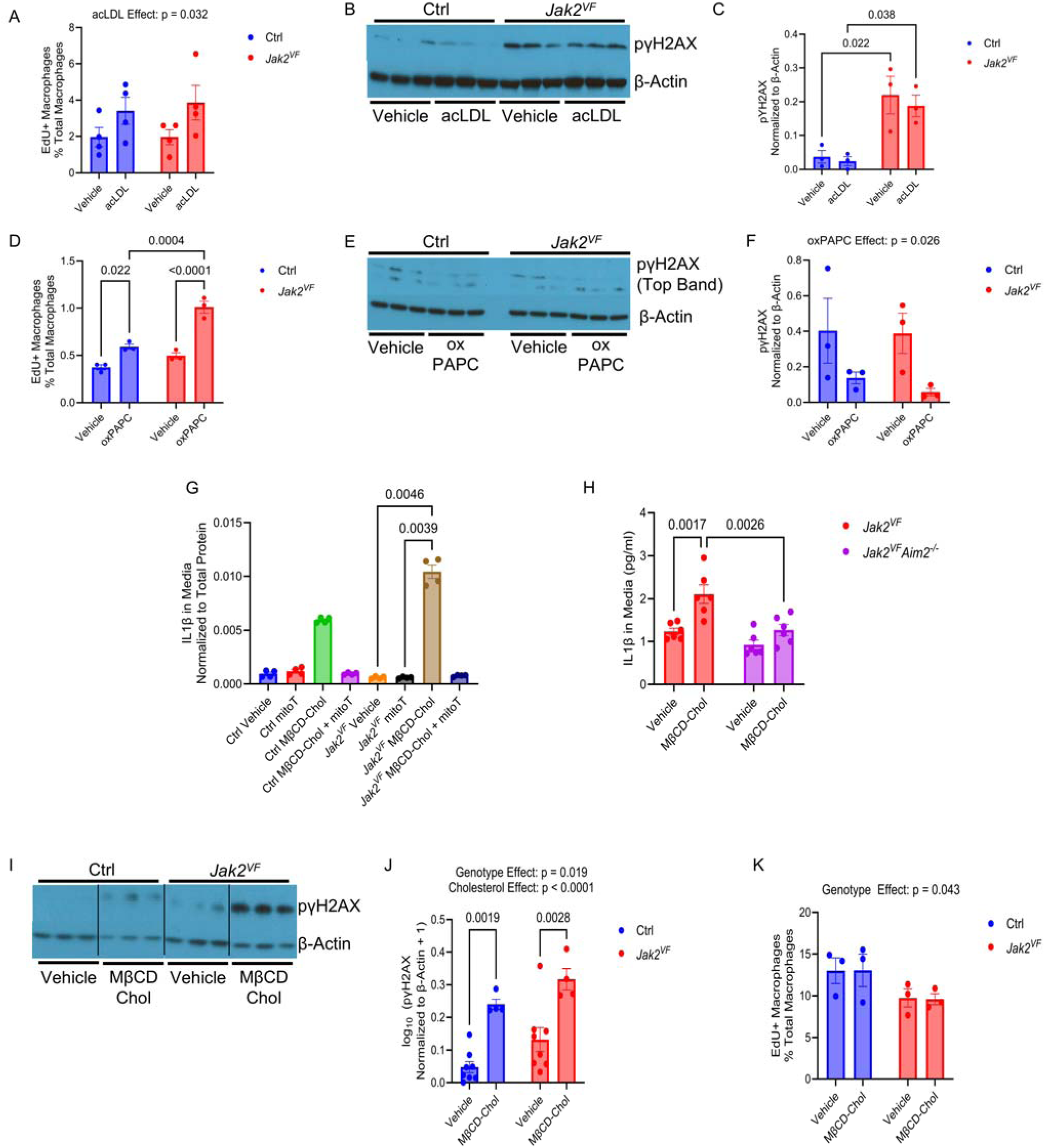
Cholesterol Increases AIM2 Inflammasome Activation and Nuclear dsDNA Breaks via Increased Mitochondrial ROS in *Jak2^VF^* Macrophages. **A.** EdU+ BMDMs as a percentage of total BMDMs after overnight incubation with acLDL (25 µg/ml). n = 4 biological replicates. *P =* 0.032 for acLDL effect. **B.** Representative immunoblot analysis of pγH2AX in BMDMs after overnight incubation with acLDL (25 µg/ml). **C.** Densitometric quantification of pγH2AX normalized to β-actin from B. Each data point represents a cell-well technical replicate from one mouse (1 biological replicate, 3 technical replicates). *P =* 0.022 (Ctrl Vehicle vs. *Jak2^VF^* Vehicle), *P* = 0.038 (Ctrl acLDL vs. *Jak2^VF^*acLDL). Statistical comparisons in this panel were performed across cell-well technical replicates from a single mouse (1 biological replicate; n = 3 wells per condition) to assess within-experiment reproducibility. **D.** EdU+ BMDMs as a percentage of total BMDMs after overnight incubation with oxPAPC (50 µg/ml). Each data point represents a cell-well technical replicate from one mouse (1 biological replicate, 3 technical replicates). P = 0.022 (Ctrl Vehicle vs. Ctrl oxPAPC), P < 0.0001 (*Jak2^VF^* Vehicle vs. *Jak2^VF^* oxPAPC), P = 0.0004 (Ctrl oxPAPC vs. *Jak2^VF^*oxPAPC). Statistical comparisons in this panel were performed across cell-well technical replicates from a single mouse (1 biological replicate; n = 3 wells per condition) to assess within-experiment reproducibility. **E.** Representative immunoblot analysis of pγH2AX in BMDMs after overnight incubation with oxPAPC (50 µg/ml). **F.** Densitometric quantification of pγH2AX normalized to β-Actin from E. Each data point represents a cell-well technical replicate from one mouse (1 biological replicate, 3 technical replicates). P = 0.026 for oxPAPC effect. Statistical comparisons in this panel were performed across cell-well technical replicates from a single mouse (1 biological replicate; n = 3 wells per condition) to assess within-experiment reproducibility. **G.** ELISA quantification of total IL1β in cell culture media of BMDMs incubated with MβCD-cholesterol (30µg/ml) alone, mitoTEMPO (10 µM) alone, or MβCD-cholesterol (30 µg/ml) in combination with mitoTEMPO (10 µM) for 4 hours after priming with LPS (100 ng/ml) for 3 hours. n = 4 cell-well technical replicates from a single mouse (1 biological replicate). Wells failing prespecified quality-control criteria were excluded (see Statistical Analysis). Statistical comparisons in this panel were performed across cell-well technical replicates from a single mouse (1 biological replicate; n = 4 wells per condition) to assess within-experiment reproducibility. **H.** ELISA quantification of total IL1β in cell culture media of BMDMs incubated with MβCD-cholesterol (30 µg/ml) for 4 hours after priming with LPS (100 ng/ml) for 3 hours. n = 6 cell-well technical replicates from a single mouse (1 biological replicate). No biological replicates were available. *P =* 0.0017 (*Jak2^VF^* Vehicle vs. *Jak2^VF^* MβCD-cholesterol), *P* = 0.0026 (*Jak2^VF^* MβCD-cholesterol vs. *Jak2^VF^Aim2*^−/-^ MβCD-cholesterol). Statistical comparisons in this panel were performed across cell-well technical replicates from a single mouse (1 biological replicate; n = 6 wells per condition) to assess within-experiment reproducibility. **I.** Representative immunoblot analysis of pγH2AX in BMDMs after a 4-hour incubation with MβCD-cholesterol (30 µg/ml). **J.** Densitometric quantification of pγH2AX normalized to β-Actin from H. n = 8 biological replicates for the vehicle groups and n = 4 biological replicates for the MβCD-cholesterol groups. *P =* 0.0019 (Ctrl Vehicle vs. Ctrl MβCD-cholesterol), *P* = 0.0028 (*Jak2^VF^* Vehicle vs. *Jak2^VF^* MβCD-cholesterol). *P* = 0.019 for genotype effect. *P* < 0.0001 for MβCD-cholesterol effect. Ctrl Vehicle data include earlier assessments of this endpoint in untreated cells, prior to the initiation of the treatment arm. All samples were processed and analyzed using the same protocol. **K.** EdU+ BMDMs as a percentage of total BMDMs after a 4-hour incubation with MβCD-cholesterol (30 µg/ml). n = 3 biological replicates. *P =* 0.043 for genotype effect. All quantifications shown as mean ± s.e.m. Two-way ANOVA with Tukey’s multiple comparisons test (A,C,D,F,H,J,K). Kruskal-Wallis test with Dunn’s multiple comparisons test (G).

## 4. Discussion

Based on prior studies (30), we hypothesized that LDL lowering alone would not prevent plaque progression in *Jak2^VF^* mice. While this proved to be the case with moderate LDL lowering, aggressive cholesterol lowering halted progression and induced beneficial remodeling changes, including decreased macrophage burden and increased lesional collagen content similarly in controls and *Jak2^VF^*MPN mice. Surprisingly, LDL lowering reversed the major cellular changes in *Jak2^VF^* macrophages that have been associated with athero-progression. This included reduced AIM2 expression and pyroptosis and increased MerTK, TREM2 and in situ efferocytosis that have all been mechanistically linked in prior studies of *Jak2^VF^*mice (29). Our prior studies have shown that inflammasome activation leads to reduced levels of MerTK and TREM2 (29), suggesting that increases in these factors during cholesterol lowering may be secondary to reduced inflammasome activation.

Prior studies have shown that LDL lowering in mice with established atherosclerosis may induce atherosclerosis regression, limitation of progression, or beneficial remodeling involving multiple mechanisms (47) including macrophage emigration (48), decreased monocyte entry (49) and a role of acquired immunity and T-regs (17). Our studies using a cleaved GasD antibody to monitor pyroptosis in lesions, indicated that reversal of inflammasome activation may also be a key mechanism underlying beneficial remodeling of plaques to a more stable phenotype, especially in settings where there is underlying increased inflammasome activation as occurs in *Tet2* and *Jak2VF* CH. Similar to macrophage proliferation being a major factor accounting for increased macrophage number during advanced atherosclerosis progression (50), suppression of proliferation may be an important mechanism decreasing macrophage burden with LDL lowering (36). Our studies suggest that reversal of the proliferative tendency of *Jak2^VF^* macrophages is also a key mechanism decreasing macrophage burden with LDL lowering in *Jak2^VF^*mice. The mechanism of proliferation could involve uptake of modified LDL by scavenger receptors into the endo-lysosomal system (44) consistent with our observations. The decreased proliferation of *Jak2VF* macrophages signifies that there are fewer inflammatory macrophages in lesions, consistent with the virtual disappearance of *Jak2^VF^* macrophages in lesions with aggressive cholesterol lowering.

The parallel decrease in macrophage proliferation and DNA damage response suggested that decreased replication induced DNA damage could be a mechanism to explain decreased AIM2 inflammasome activation and pyroptosis. However, mechanistic studies in *Jak2^VF^*BMDMs indicated that macrophage proliferation could be clearly dissociated from DNA damage responses and AIM2 inflammasome activation that were rather dependent on mitochondrial ROS generation and DNA damage as suggested by earlier studies (46,51). The increased DNA damage and inflammasome activation in *Jak2^VF^* macrophages compared to controls likely reflects increased glycolysis, mitochondrial ROS formation and oxidative mitochondrial damage (30,52). However, the decreased *Jak2^VF^* macrophage burden, as well as decreased AIM2 expression and decreased cholesterol-induced AIM2 inflammasome activation together explain why effective LDL lowering was sufficient to reverse the adverse effects of *Jak2^VF^* on atherosclerosis.

Paradoxically, despite the overall reduction in macrophage burden, during resolution there may also be a relative increase in a proliferative set of efferocytotic macrophages that have increased c-myc expression and contribute to plaque stability (13,14). In addition to a potential role in cell proliferation, c-myc increases Akt signaling and cell survival (53,54). Previously, we observed a 4-fold increase in TREM2^Hi^ macrophages induced by both LDL lowering and *Jak2^VF^*inactivation in *Jak2^VF^* CH mice (16). In the present study, we found that LDL lowering alone led to a marked increase in TREM2 staining in macrophages, as well as elevated c-myc expression specifically within TREM2^Hi^ macrophages, in both control and mutant mice. This contrasts with the decreased proliferation and DNA damage response found specifically in the *Jak2^VF^* macrophages. Our findings together suggest that there is an increase in TREM2^Hi^, c-myc and MerTK expressing macrophages induced by LDL lowering either through increased proliferation or increased survival of these subsets that then have beneficial effects on efferocytosis and plaque stability. Given that inflammasome activation and IL-1 release have been linked to reduced MerTK and TREM2 as a result of a disintegrin and metalloproteinase 17 (ADAM17)-mediated cleavage (39,55), the increase in this pro-resolving macrophage population may in part be secondary to reduced cholesterol-induced inflammasome activation.

LDL lowering guidelines have recommended progressively lower targets based on clinical trial evidence. Current recommendations suggest an LDL cholesterol target below 70 mg/dl for patients with established ACVD or in certain high-risk groups (56). Statin use has been associated with improved outcomes in human MPN patients and was associated with a 37% reduction in the risk of thrombosis in patients who had polycythemia vera or essential thrombocythemia as well as a 22% reduction in all-cause mortality (57). While data from mice should be extrapolated to humans with caution, the AIM2 inflammasome seems to increase ACVD risk in both mice and humans carrying the *Jak2^VF^* mutation (31). The present preclinical study adds mechanistic findings to support the idea that aggressive LDL-C lowering in *Jak2^VF^* MPN patients would reduce ACVD risk.

## Abbreviations

AAV8: Adeno-Associated Virus serotype 8 vector
acLDL: acetylated LDL
ACVD: atherosclerotic cardiovascular disease
ADAM17: a disintegrin and metalloproteinase 17
AIM2: absent in melanoma 2
BMDM: bone marrow-derived macrophages
BMT: bone marrow transplantation
CH: clonal hematopoiesis
CHIP: clonal hematopoiesis of indeterminate potential
FIJI: FIJI Is Just ImageJ
GasD: gasdermin D
hDAD: helper-dependent adenoviral vector
HSC: hematopoietic stem cell
IL-1: interleukin-1
IL-1β: interleukin-1β
*Jak2^VF^*: *Jak2^V617F^*
MerTK: mer proto-oncogene tyrosine kinase
MitoTEMPO: mitochondria-targeted TEMPO (a triphenylphosphonium-conjugated nitroxide)
MPN: myeloproliferative neoplasms
MβCD: methyl-β-cyclodextrin
oxPAPC: oxidized 1-palmitoyl-2-arachidonoyl-sn-glycero-3-phosphocholine
pIpC: polyinosinic:polycytidylic acid
pγH2AX: phosphorylated histone H2A.X
TREM2: triggering receptor expressed on myeloid cells 2

## Acknowledgements

The authors thank Wenli Liu for assistance and support during the course of this study. This study used the Confocal and Specialized Microscopy Shared Resource of the Herbert Irving Comprehensive Cancer Center at Columbia University, funded in part through NIH/NCI Cancer Center Support Grant P30CA013696. Additionally, this study used the resources of the Herbert Irving Comprehensive Cancer Center Flow Cytometry Shared Resources funded in part through Center Grant P30CA013696.

## Data Availability

The datasets generated for this study are available from the corresponding author upon reasonable request.

