## Supplementary Materials for "Aggressive Cholesterol Lowering Normalizes Atherosclerosis Regression in *Jak2^V617F^* Mice"

**Supplemental Material for Aggressive Cholesterol Lowering Normalizes Atherosclerosis Regression in *Jak2^V617F^* Mice**

**Supplemental Figures**

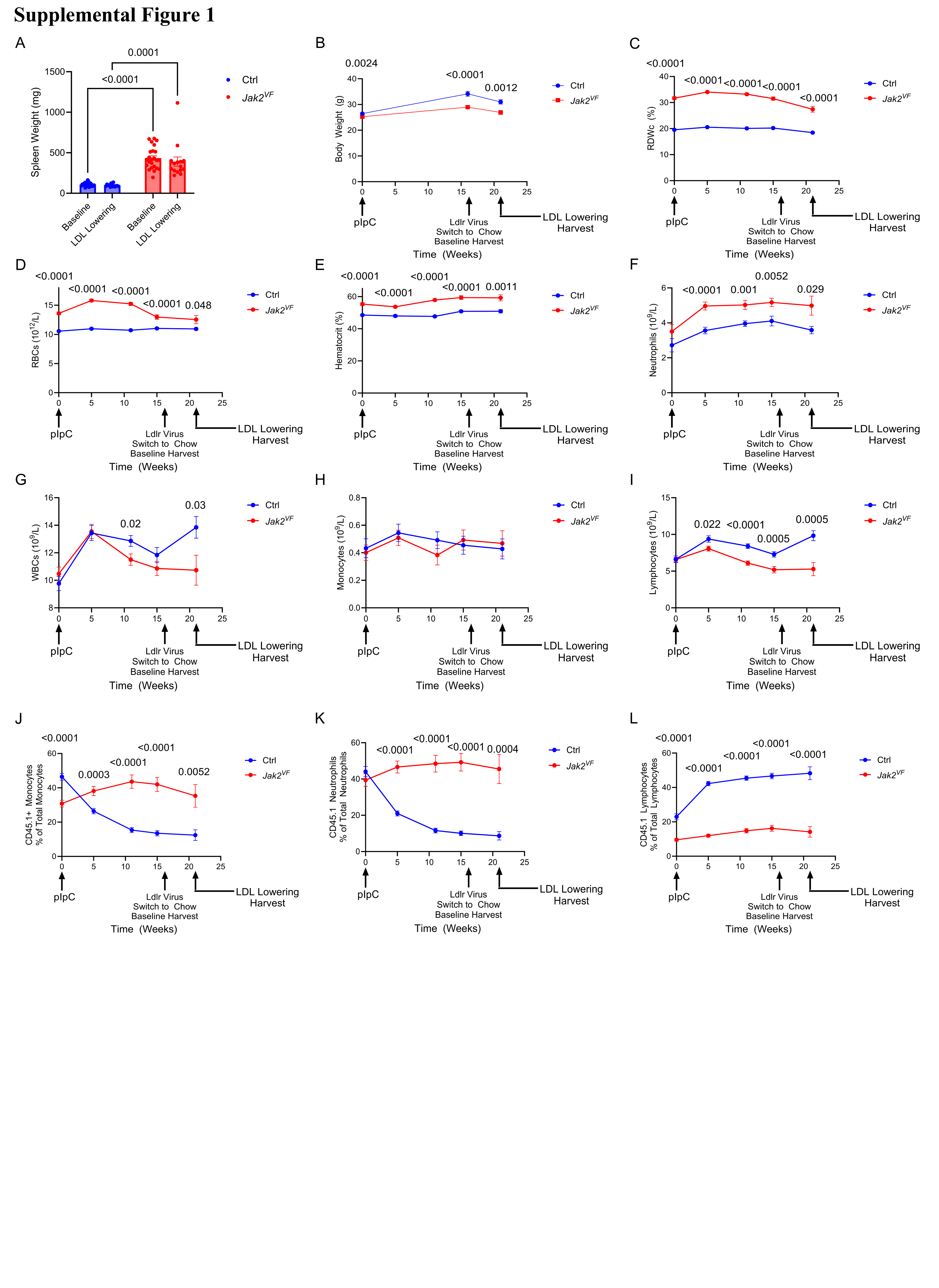

**Supplemental Figure 1.** ***Jak2^VF^* Burden and Blood Cell Counts in Moderate Cholesterol Lowering Mx1-Cre Study. A.** Spleen weight, n = 14-24. *P* < 0.0001 (Ctrl Baseline vs *Jak2^VF^* Baseline), P = 0.0001 (Ctrl LDL Lowering vs *Jak2^VF^* LDL Lowering). **B.** Body weight *P* = 0.0024, <0.0001, 0.0012 (Ctrl vs *Jak2^VF^* at weeks 0, 16, 21 respectively). **C.** RDWc. *P* <0.0001 (Ctrl vs *Jak2^VF^* for all timepoints). **D.** RBCs. *P* <0.0001, 0.048 (Ctrl vs *Jak2^VF^* at weeks 0-15 and 21 respectively). **E.** Hematocrit. *P* <0.0001, 0.0011 (Ctrl vs *Jak2^VF^* at weeks 0-15 and 21 respectively). **F.** Neutrophils. *P* < 0.0001, 0.001, 0.0052, 0.029 (Ctrl vs *Jak2^VF^* at weeks 5, 11, 15, 21 respectively). **G.** WBCs. *P* = 0.02, 0.03 (Ctrl vs *Jak2^VF^* at weeks 11, 21 respectively). **H.** Monocytes. **I.** Lymphocytes. *P* = 0.022, <0.0001, 0.0005, 0.0005 (Ctrl vs *Jak2^VF^* at weeks 5, 11, 15, 21 respectively). **J.** Percentage of total monocytes positive for CD45.1. *P* <0.0001, 0.0003, <0.0001, <0.0001, 0.0052 (Ctrl vs *Jak2^VF^* at weeks 0, 5, 11, 15, 21 respectively). **K.** Percentage of total neutrophils positive for CD45.1. *P* <0.0001, <0.0001, <0.0001, 0.0004 (Ctrl vs *Jak2^VF^* at weeks 5, 11, 15, 21 respectively). **L.** Percentage of total lymphocytes positive for CD45.1. *P* <0.0001 (Ctrl vs *Jak2^VF^* at all timepoints). All quantifications shown as mean ± s.e.m. For B, n = 15-43. For C-I, n = 11-52. For J-L, n = 15-40. Kruskal-Wallis test with Dunn’s multiple comparisons test (A). Two-way ANOVA with the Geisser-Greenhouse correction for sphericity and Tukey’s multiple comparisons test (B-L).

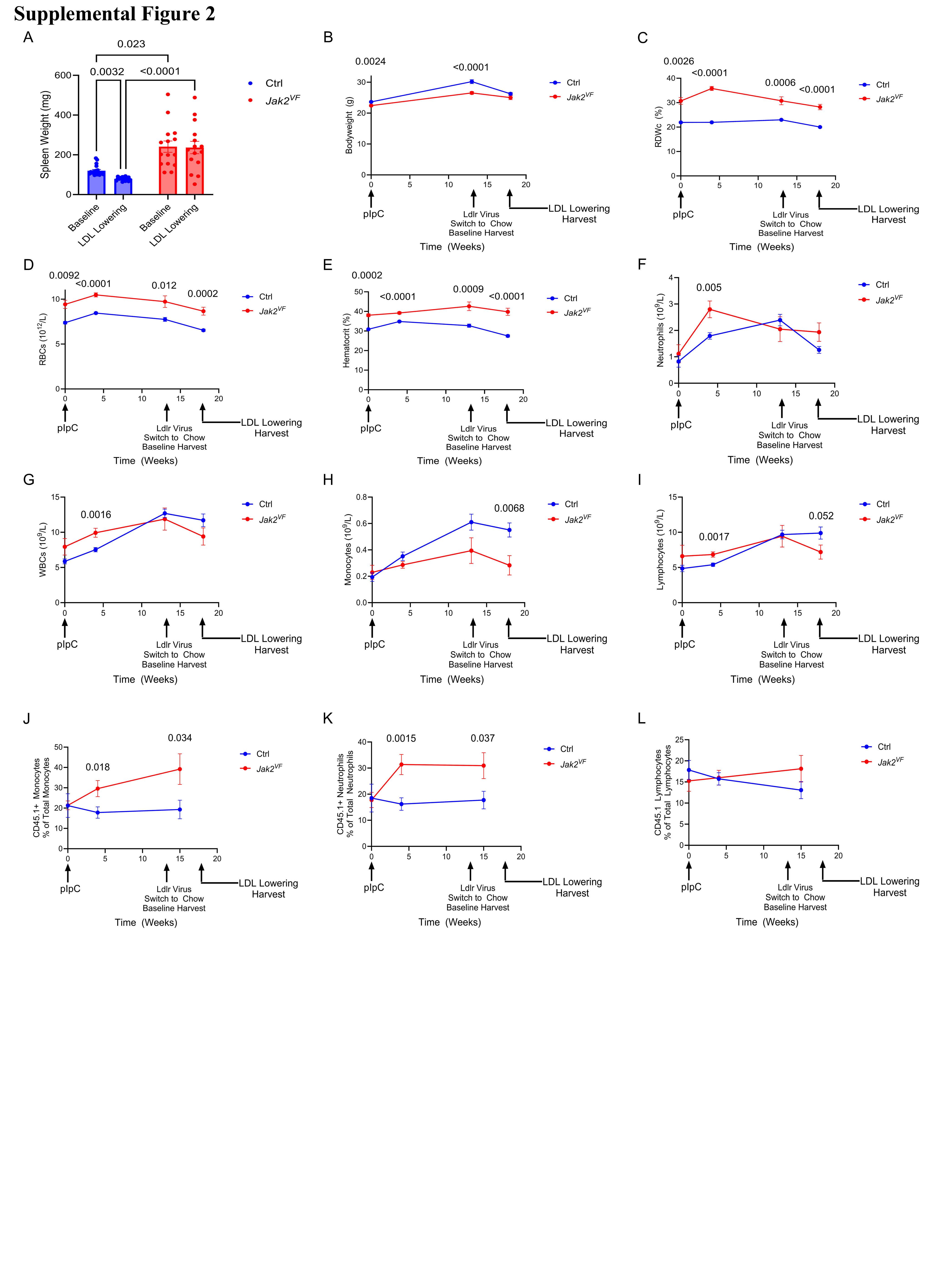

**Supplemental Figure 2.** ***Jak2^VF^* Burden and Blood Cell Counts in Aggressive Cholesterol Lowering Mx1-Cre Study. A.** Spleen weight, n = 14-20. *P* = 0.023 (Ctrl Baseline vs *Jak2^VF^* Baseline), P < 0.0001 (Ctrl LDL Lowering vs *Jak2^VF^* LDL Lowering), *P* = 0.0032 (Ctrl Baseline vs LDL Lowering **B.** Body weight. *P* = 0.0024, <0.0001 (Ctrl vs *Jak2^VF^* at weeks 0, 13 respectively). **C.** RDWc. *P* = 0.0026, <0.0001, 0.0006, <0.0001 (Ctrl vs *Jak2^VF^* at weeks 0, 4, 13, 18 respectively). **D.** RBCs. *P* = 0.0092, <0.0001, 0.012, 0.0002 (Ctrl vs *Jak2^VF^* at weeks 0, 4, 13, 18 respectively). **E.** Hematocrit. *P* = 0.0002, <0.0001, 0.0009, <0.0001 (Ctrl vs *Jak2^VF^* at weeks 0, 4, 13, 18 respectively). **F.** Neutrophils. *P* = 0.005 (Ctrl vs *Jak2^VF^* at week 4). **G.** WBCs. *P* = 0.0016 (Ctrl vs *Jak2^VF^* at week 4). **H.** Monocytes. *P* = 0.0068 (Ctrl vs *Jak2^VF^* at week 18). **I.** Lymphocytes. *P* = 0.0017, 0.052 (Ctrl vs *Jak2^VF^* at weeks 4, 18 respectively). **J.** Percentage of total monocytes positive for CD45.1. *P* = 0.018, 0.034 (Ctrl vs *Jak2^VF^* at weeks 4, 15 respectively). **K.** Percentage of total neutrophils positive for CD45.1. *P* = 0.0015, 0.037 (Ctrl vs *Jak2^VF^* at weeks 4, 15 respectively). **L.** Percentage of total lymphocytes positive for CD45.1. All quantifications shown as mean ± s.e.m. For B, n = 18-38. For C-I, n = 5 for week 0, n = 15-36 for weeks 4-18. For J-L, n = 5 for week 0, n = 16-39 for weeks 4-15. Kruskal-Wallis test with Dunn’s multiple comparisons test (A). Two-way ANOVA with the Geisser-Greenhouse correction for sphericity and Tukey’s multiple comparisons test (B-L).

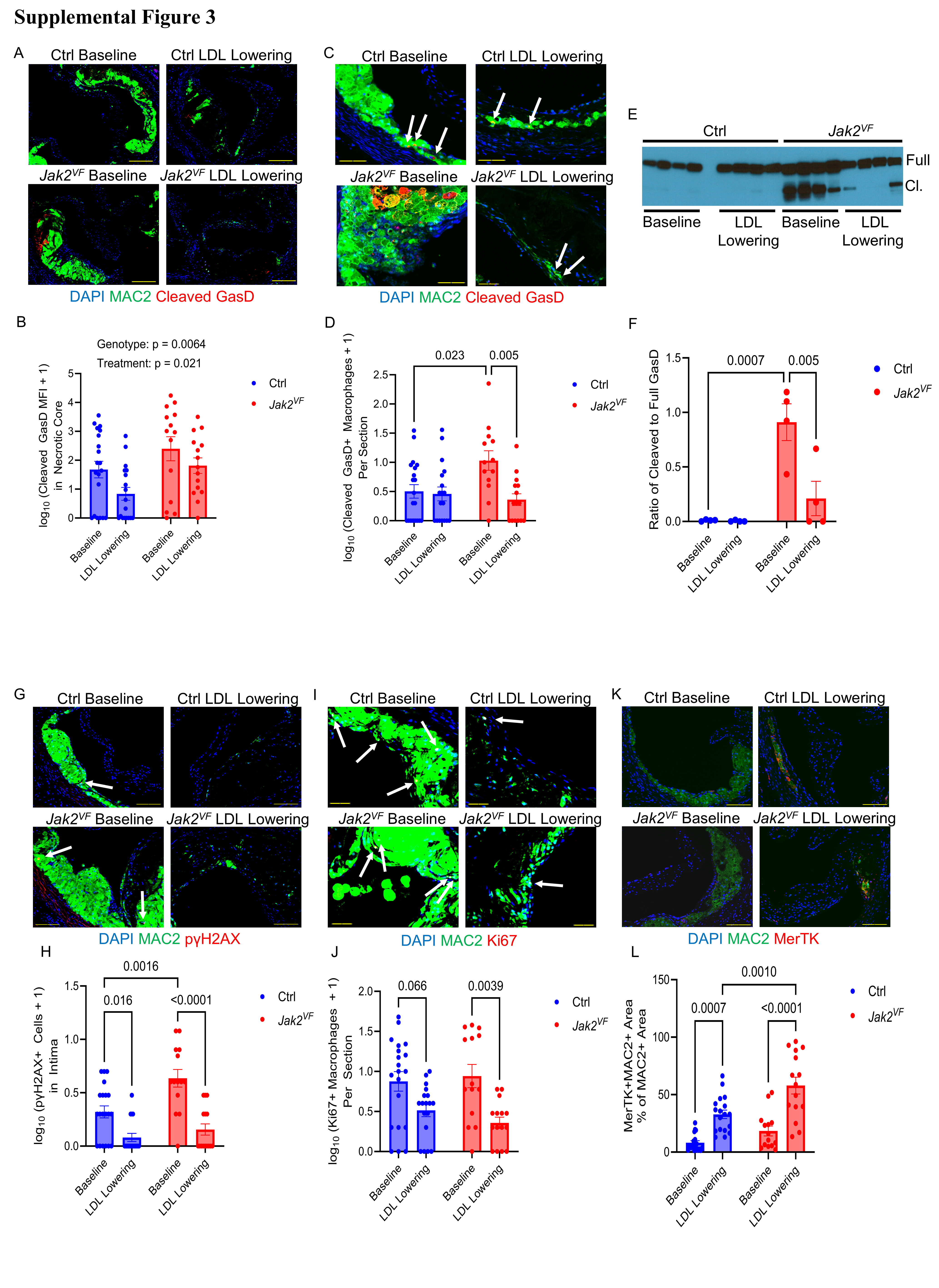

**Supplemental Figure 3.** **Aggressive Cholesterol Lowering More Strongly Decreases Macrophage Pyroptosis and DNA Damage while Similarly Increasing Macrophage MerTK in *Jak2^VF^* MPN Lesions. A.** Images of aortic root lesions stained for MAC2 (Green), Cleaved GasD (Red), and DAPI (Blue). Scale bar, 247 µm. **B.** Log_10_ transformed cleaved GasD MFI in the necrotic core with the addition of constant 1, n = 14-20. *P =* 0.0064 for genotype effect and *P* = 0.021 for treatment effect by two-way ANOVA. **C.** Images of aortic root lesions stained for MAC2 (Green), Cleaved GasD (Red), and DAPI (Blue). Scale bar, 96 µm. White arrows, cleaved GasD+ macrophages. **D.** Log_10_ transformed cleaved GasD positive macrophages per section with the addition of constant 1, n = 13-20. *P* = 0.023 (Ctrl Baseline vs *Jak2^VF^* Baseline), *P* = 0.005 (*Jak2^VF^* Baseline vs LDL Lowering). **E.** Representative immunoblot analysis of full-length and cleaved GasD in CD11b+ splenocytes. Full, full-length GasD, Cl., cleaved GasD. **F.** Densitometric quantification of the ratio of cleaved GasD to full-length GasD from E. n = 4 biological replicates. *P* = 0.0007 (Ctrl Baseline vs *Jak2^VF^* Baseline), *P* = 0.005 (*Jak2^VF^* Baseline vs LDL Lowering). **G.** Images of aortic root lesions stained for MAC2 (Green), pγH2AX (Red), and DAPI (Blue). Scale bar, 100 µm. White arrows, pγH2AX positive cells. **H.** Log_10_ transformed pγH2AX positive macrophages per section with the addition of constant 1, n = 14-20. *P* = 0.016 (Ctrl Baseline vs LDL Lowering), *P* = 0.0016 (Ctrl Baseline vs *Jak2^VF^* Baseline), *P* < 0.0001 (*Jak2^VF^* Baseline vs LDL Lowering). **I.** Images of aortic root lesions stained for MAC2 (Green), Ki67 (Red), and DAPI (Blue). Scale bar, 40 µm. White arrows, Ki67 positive macrophages. **J.** Log_10_ transformed Ki67 positive macrophages per section with the addition of constant 1, n = 14-20. *P* = 0.066 (Ctrl Baseline vs LDL Lowering), *P* = 0.0039 (*Jak2^VF^* Baseline vs LDL Lowering). **K.** Images of aortic root lesions stained for MAC2 (Green), MerTK (Red), and DAPI (Blue). Scale bar, 100 µm. **L.** Percentage of MAC2 positive area double positive for MerTK and MAC2, n = 14-18. *P* = 0.0007 (Ctrl Baseline vs LDL Lowering), *P* = 0.0010 (Ctrl LDL Lowering vs *Jak2^VF^* LDL Lowering), *P* < 0.0001 (*Jak2^VF^* Baseline vs LDL Lowering). All quantifications shown as mean ± s.e.m. Two-way ANOVA with Tukey’s multiple comparisons test (B,D,F,H,J,L).

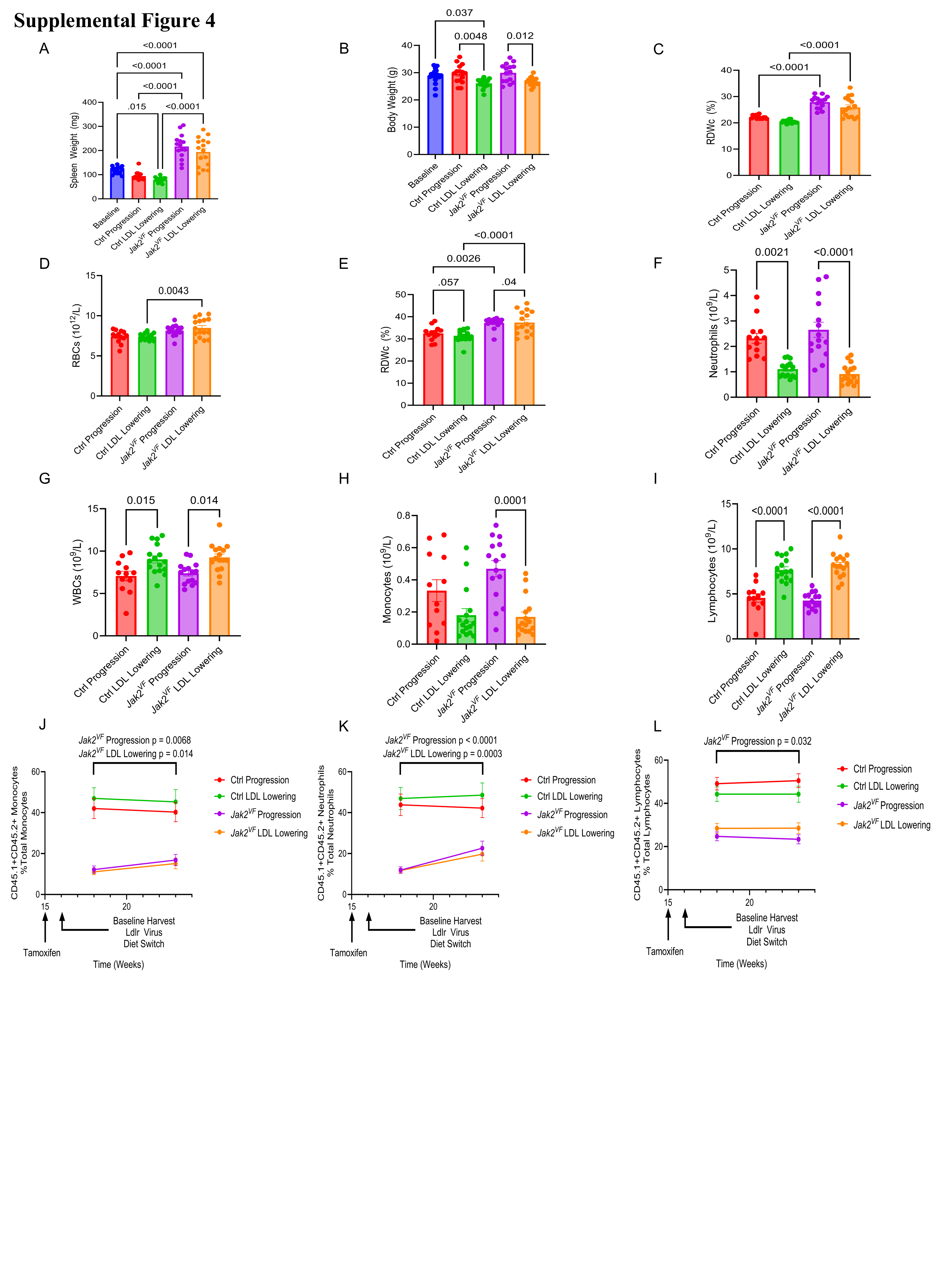

**Supplemental Figure 4.** ***Jak2^VF^* Burden and Blood Cell Counts in Aggressive Cholesterol Lowering Scl-Cre Study. A.** Spleen weight, n = 14-16. *P* = 0.015 (Baseline vs Ctrl LDL Lowering), *P* < 0.0001 (Baseline vs *Jak2^VF^* Progression; Baseline vs *Jak2^VF^* LDL Lowering; Ctrl Progression vs *Jak2^VF^* Progression; Ctrl LDL Lowering vs *Jak2^VF^* LDL Lowering). **B.** Body weight, n = 16-17. *P* = 0.037 (Baseline vs Ctrl LDL Lowering), *P* = 0.0048 (Ctrl Progression vs Ctrl LDL Lowering), *P* = 0.012 (*Jak2^VF^* Progression vs *Jak2^VF^* LDL Lowering). **C.** RDWc, n = 13-16. *P* < 0.0001 (Ctrl Progression vs *Jak2^VF^* Progression; Ctrl LDL Lowering vs *Jak2^VF^* LDL Lowering). **D.** RBCs, n = 13-16. *P* = 0.0043 (Ctrl LDL Lowering vs *Jak2^VF^* LDL Lowering). **E.** Hematocrit, n = 13-16. *P* = 0.0007 (Ctrl Progression vs *Jak2^VF^* Progression), *P* = 0.0001 Ctrl LDL Lowering vs *Jak2^VF^* LDL Lowering). **F.** Neutrophils, n = 13-16. *P* = 0.0021 (Ctrl Progression vs Ctrl LDL Lowering), *P* < 0.0001 (*Jak2^VF^* Progression vs *Jak2^VF^* LDL Lowering). **G.** WBCs, n = 12-16. *P* = 0.015 (Ctrl Progression vs Ctrl LDL Lowering), *P* = 0.014 (*Jak2^VF^* Progression vs *Jak2^VF^* LDL Lowering). **H.** Monocytes, n = 12-16. *P* = 0.0001 (*Jak2^VF^* Progression vs *Jak2^VF^* LDL Lowering). **I.** Lymphocytes, n = 12-16. *P* < 0.0001 (Ctrl Progression vs Ctrl LDL Lowering), *P* < 0.0001 (*Jak2^VF^* Progression vs *Jak2^VF^* LDL Lowering). **J.** Percentage of total monocytes double positive for CD45.1 and CD45.2, n = 29-31. *P* = 0.0068 (*Jak2^VF^* Progression 18 weeks vs 23 weeks), *P* = 0.014 (*Jak2^VF^* LDL Lowering 18 weeks vs 23 weeks). **K.** Percentage of total neutrophils double positive for CD45.1 and CD45.2, n = 29-31. *P* < 0.0001 (*Jak2^VF^* Progression 18 weeks vs 23 weeks), *P* = 0.0003 (*Jak2^VF^* LDL Lowering 18 weeks vs 23 weeks). **L.** Percentage of total lymphocytes positive double positive for CD45.1 and CD45.2, n = 29-31. *P* = 0.032 (*Jak2^VF^* Progression 18 weeks vs 23 weeks). All quantifications shown as mean ± s.e.m. B-I were taken at time of harvest. One-way ANOVA with Holm-Sidak’s multiple comparisons test (A), Two-way ANOVA with Tukey’s multiple comparisons test (B,C,D,E,G,H,I), Kruskal-Wallis test with Dunn’s multiple comparisons test (F). Two-way ANOVA with the Geisser-Greenhouse correction for sphericity and Tukey’s multiple comparisons test (J-L).

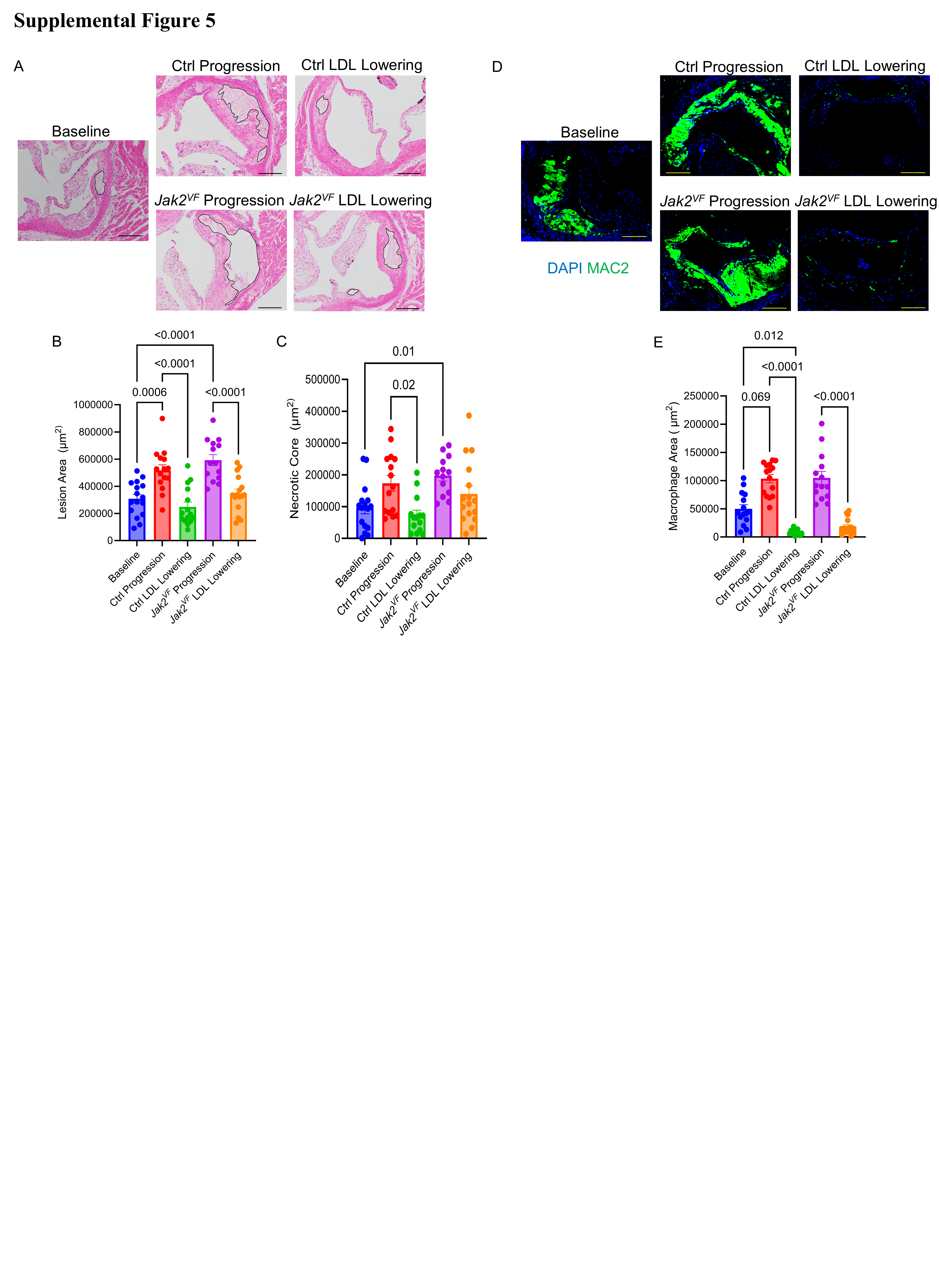

**Supplemental Figure 5.** **Aggressive Cholesterol Lowering Normalizes Regression in *Jak2^VF^* Mice with Mutation Activated After Established Atherosclerosis. A.** H&E images of aortic root lesions. Black lines, necrotic core. Scale bar, 200 µm. **B.** Lesion area, n = 14-16. *P* = 0.0006 (Baseline vs Ctrl Progression), *P* < 0.0001 (Baseline vs *Jak2^VF^* Progression; Ctrl Progression vs Ctrl LDL Lowering; *Jak2^VF^* Progression vs *Jak2^VF^* LDL Lowering). **C.** Necrotic core area, n = 13-16. *P* = 0.01 (Baseline vs *Jak2^VF^* Progression), *P* = 0.02 (Ctrl Progression vs Ctrl LDL Lowering). **D.** Images of aortic root lesions stained for MAC2 (Green) and DAPI (Blue). Scale bar, 159 µm. **E.** Macrophage area, n = 14-16. *P* = 0.069 (Baseline vs Ctrl Progression), *P* = 0.012 (Baseline vs Ctrl LDL Lowering), *P* < 0.0001 (Ctrl Progression vs Ctrl LDL Lowering; *Jak2^VF^* Progression vs *Jak2^VF^* LDL Lowering). All quantifications shown as mean ± s.e.m. One-way ANOVA with Holm-Sidak’s multiple comparisons test (B) Kruskal-Wallis test with Dunn’s multiple comparisons test (C,E).

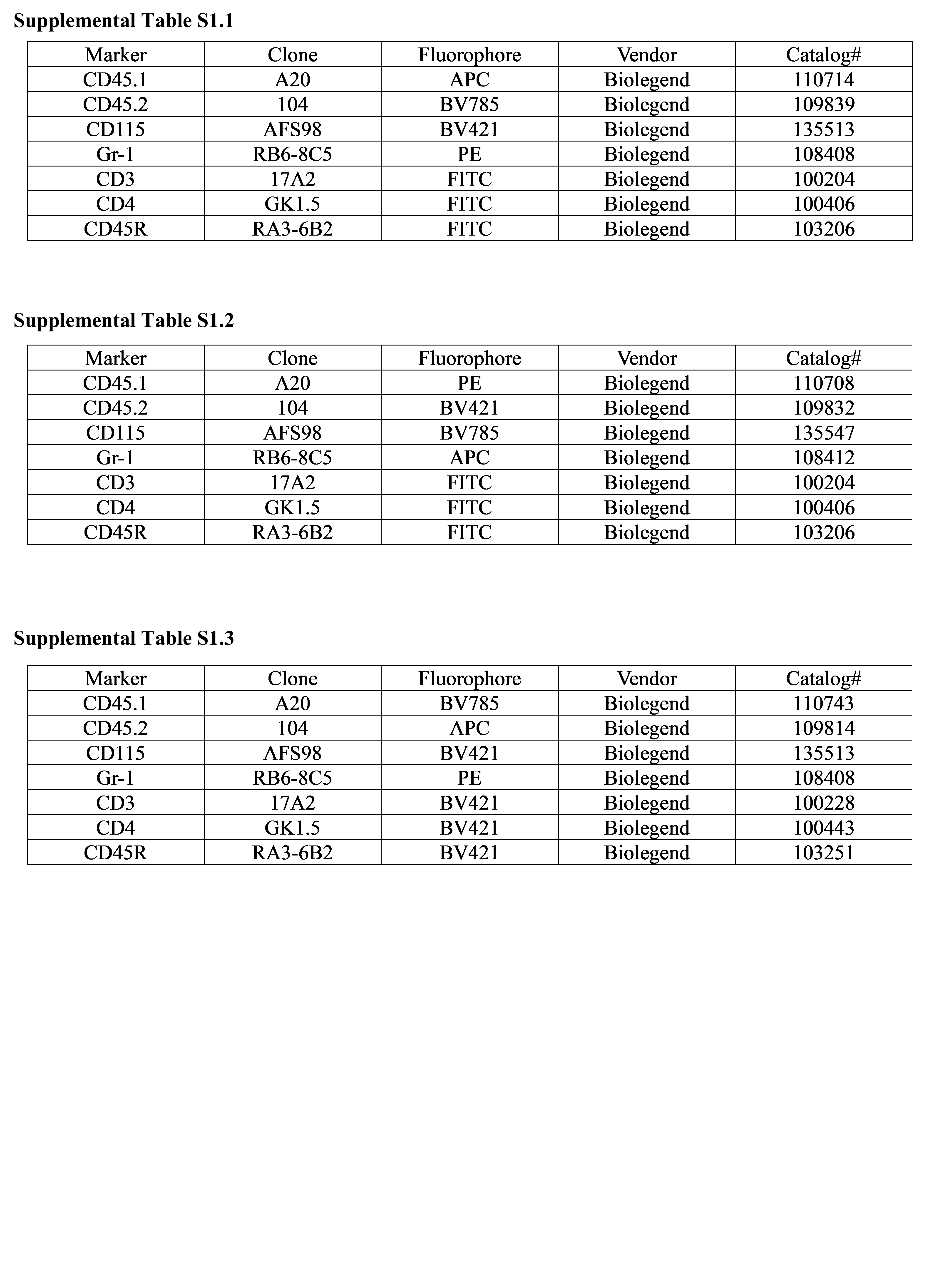

**Supplemental Table S1.** **Antibodies Used for Flow Cytometric Analysis of Blood Leukocytes**
